# Fiber alignment in 3D collagen networks as a biophysical marker for cell contractility

**DOI:** 10.1101/2023.06.28.546896

**Authors:** David Böhringer, Andreas Bauer, Ivana Moravec, Lars Bischof, Delf Kah, Christoph Mark, Thomas J. Grundy, Ekkehard Görlach, Geraldine M O’Neill, Silvia Budday, Pamela Strissel, Reiner Strick, Andrea Malandrino, Richard Gerum, Michael Mak, Martin Rausch, Ben Fabry

## Abstract

Cells cultured in 3D fibrous biopolymer matrices exert traction forces on their environment that induce deformations and remodeling of the fiber network. By measuring these deformations, the traction forces can be reconstructed if the mechanical properties of the matrix and the force-free matrix configuration are known. These requirements severely limit the applicability of traction force reconstruction in practice. In this study, we test whether force-induced matrix remodeling can instead be used as a proxy for cellular traction forces. We measure the traction forces of hepatic stellate cells and different glioblastoma cell lines and quantify matrix remodeling by measuring the fiber orientation and fiber density around these cells. In agreement with simulated fiber networks, we demonstrate that changes in local fiber orientation and density are directly related to cell forces. By resolving Rho-kinase (ROCK) Inhibitor-induced changes of traction forces and fiber alignment and density in hepatic stellate cells, we show that the method is suitable for drug screening assays. We conclude that differences in local fiber orientation and density, which are easily measurable, can be used as a qualitative proxy for changes in traction forces. The method is available as an open-source Python package with a graphical user interface.

## Introduction

Actin and myosin are two highly abundant proteins in most cells, which enables them to generate forces for contraction (1–3) and a wide range of other essential biological processes including cell division (4, 5), mechano-sensing (6, 7), migration (8, 9), or wound closure (10, 11). Pathological processes such as inflammatory diseases (12), fibrosis (13–15), and epithelial-to-mesenchymal transition in cancer (16) are often accompanied by changes in cellular force generation.

To study cellular force generation in vitro, different qualitative and quantitative methods have been developed. For traction force measurements under 2D cell culture conditions, cells can be seeded on a thin, flexible silicone membrane. Cellular forces cause the membrane to wrinkle, from which the location and magnitude of cell tractions can be estimated (17–19). Quantitative and locally highly resolved 2D maps of traction forces are obtained by plating cells on flat, elastic polyacrylamide hydrogels, with beads embedded near the gel surface. From the bead movements that arise when cells are trypsinized or when cell forces are relaxed with pharmacological agents (e.g using cytochalasin D), the cell forces can be reconstructed (20–22).

For traction force measurements under 3D cell culture conditions, the shrinkage of a floating collagen gel with embedded cells can be measured over time. The speed and extent of collagen gel shrinkage gives a rough qualitative measure of the overall cellular force generation (23).

It is also possible to obtain a local map of traction forces generated by cells in a 3D collagen gel (24, 25). However, 3D traction force measurements (3D TFM) in highly non-linear biopolymer hydrogels, such as collagen gels, are technically demanding, as a relatively large volume of the matrix surrounding a cell must be repeatedly imaged using confocal microscopy. Moreover, the mechanical behaviour of the extracellular matrix must be measured and described by a suitable material model. This in itself is challenging: when cells exert traction forces on the fibers of a 3D network (e.g. collagen or fibrin), some fibers buckle, others align in force direction (26, 27), and this in turn may cause a local softening or stiffening of the matrix (25, 28). In addition, 3D force evaluation requires significant computational effort to solve an inverse, mathematically ill-posed nonlinear problem using finite element analysis. Due to their complexity, 3D force measurements are rarely performed.

In this study, we developed a method to quantify the degree of local fiber orientation and density around contractile cells in 3D fiber networks. Based on computational simulations of fiber networks as well as cell experiments in collagen networks, we demonstrate that higher fiber alignment and density correlate with higher cell contractility. We exploit this correlation to measure and interpret fiber orientation and density around contractile cells as a proxy for cellular forces in 3D collagen networks. Compared to regular 3D traction force measurements, this method requires 3D imaging of smaller volumes, no additional imaging after drug-induced relaxation, no knowledge of the rheological properties of the extracellular matrix, and much less computational power for data analysis (Fig. 1). The method can be applied as a drug-screening assay and is available as an open source Python package (29).

**Fig. 1.**
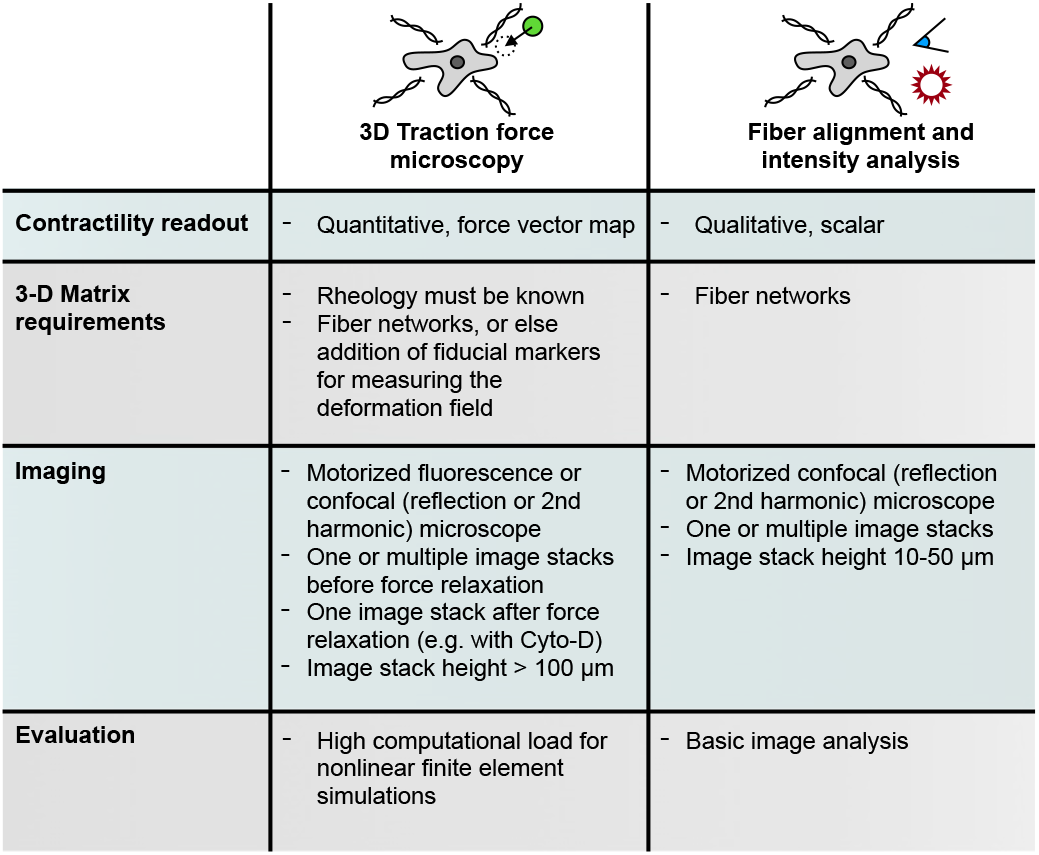
Comparison of traction force microscopy vs. quantification of matrix remodelling by fiber alignment and fiber intensity.

## Results

### Quantification and validation of fiber alignment

In this study, we embed calcein-stained hepatic stellate cells in a 3D collagen type I hydrogel and acquire image stacks of the fiber network around individual cells using second harmonic generation microscopy or confocal reflection microscopy (Fig. 2a, SI Fig. 1,3, SI Video 13,14). The direction of the collagen fibers (Fig. 2b) is determined using structure tensor analysis (30–35) as explained in Methods. We then compute the angular deviation between the fiber direction and the vector pointing to the cell center, and transform the angular deviation into an orientation value (Eq. 6) that ranges from -1 (fibers are oriented perpendicular to the center-line) to +1 (fibers are oriented parallel to the center-line (Fig. 2c).

**Fig. 2.**
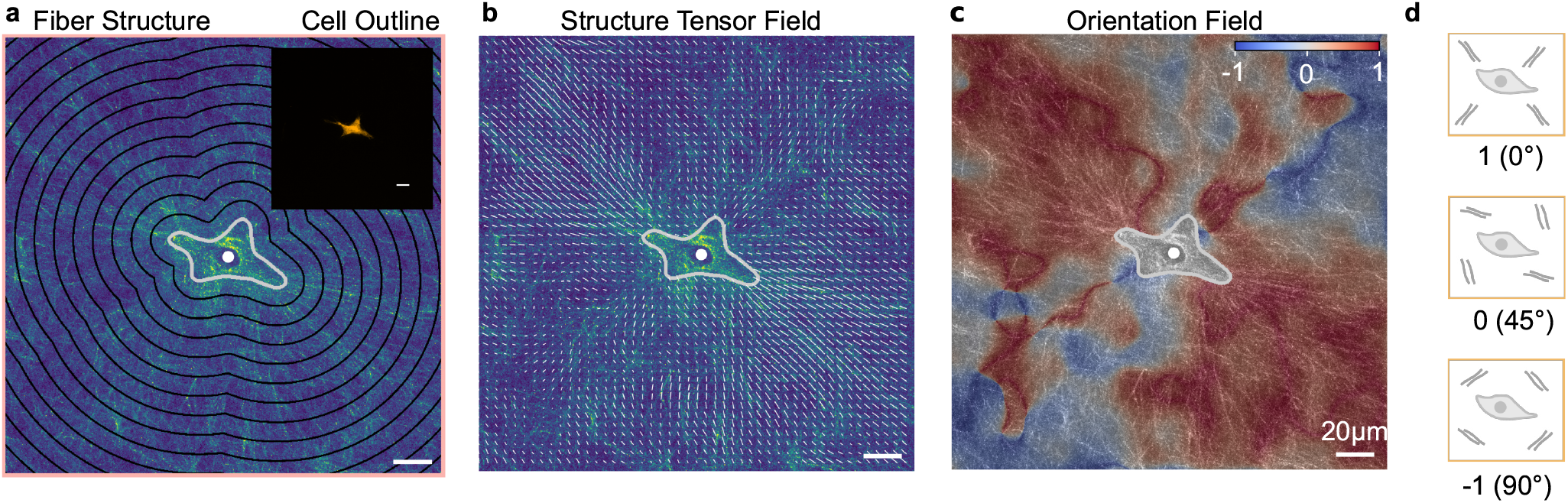
Analysis of fiber alignment: **a**, Maximum intensity projection of the collagen fiber structure around individual hepatic stellate cells imaged using second harmonic generation microscopy. The cell outline is determined from calcein-labeled images (inset). Black lines indicate lines of equal distances from the cell outline (“shells”). **b**, Fiber orientation vectors calculated using structure tensor analysis (Eq. 2). Arrow length is scaled by the coherence (anisotropy) of the structure (longer arrows indicate higher degree of alignment) (Eq. 5). **c**, Orientation field showing the magnitude of the angle between fiber orientation vectors and the vector pointing to the center of the segmented cell, normalized according to Eq. 6, as illustrated in (**d**). **d**, Fibers perpendicular to the vector pointing to the center of the segmented cell have an orientation of -1, fibers parallel to the cell vector have an orientation of +1, and fibers with an angle of ±45°have an orientation of 0.

**Fig. 3.**
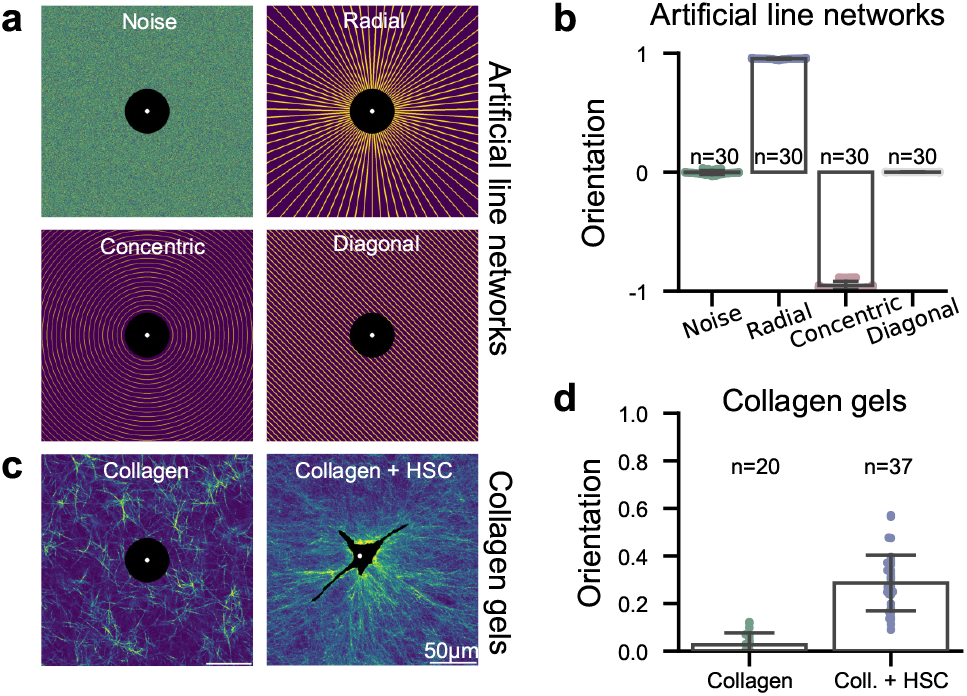
Validation of fiber orientation analysis: **a**, Fiber networks are simulated around circular non-contractile inclusions as Gaussian noise (representing dense, random fibers below the microscope’s resolution limit), or as diagonal, radial, or concentric lines. **b**, The resulting mean values of fiber orientation corresponding to the geometries of the simulated fiber networks **c**, A non-contractile inclusion is placed at the center of a confocal image of a 1.0 mg/m collagen gel polymerized at room temperature (left). Fiber orientation is compared to collagen gels containing contractile hepatic stellate cells after four days of culture (right). **d**, The mean fiber orientation confirms that collagen gels show random fiber orientation and that contractile cells induce fiber alignment (mean±sd, n = number of analyzed images).

To validate the analysis, we generate artificial images of networks with radially or concentrically aligned fibers of varying thickness and density (Fig. 3a). Additionally, we simulate diagonal line networks and images containing only Gaussian noise. The cell is represented by a sphere at the image center. As expected, for radially oriented line networks our method yields an orientation close to unity. Gaussian noise and diagonal lines both show an orientation close to 0, and concentric line networks show an orientation close to -1 (Fig. 3b).

To verify that fiber orientation is indeed a result of cell contractility and not an intrinsic property of the extracellular matrix (ECM), we analyze confocal reflection images of collagen gels containing no cells (Fig. 3c). These fiber networks show a near random orientation (0.027±0.050; mean±sd). In contrast, fibers in collagen gels (1 mg/ml) containing hepatic stellate cells (HSCs) after four days of culture are aligned towards the cell center, with a pronounced increase in mean orientation (0.286±0.117; Fig. 3d).

### Fiber networks around contractile cells

To investigate how contractile forces generated by cells give rise to spatial profiles of the fiber network orientation values, we simulate different cell-ECM interactions. First, we generate a disordered network of elastic fibers with no preferred orientation, mimicking an unloaded ECM (Fig. 4a).

**Fig. 4.**
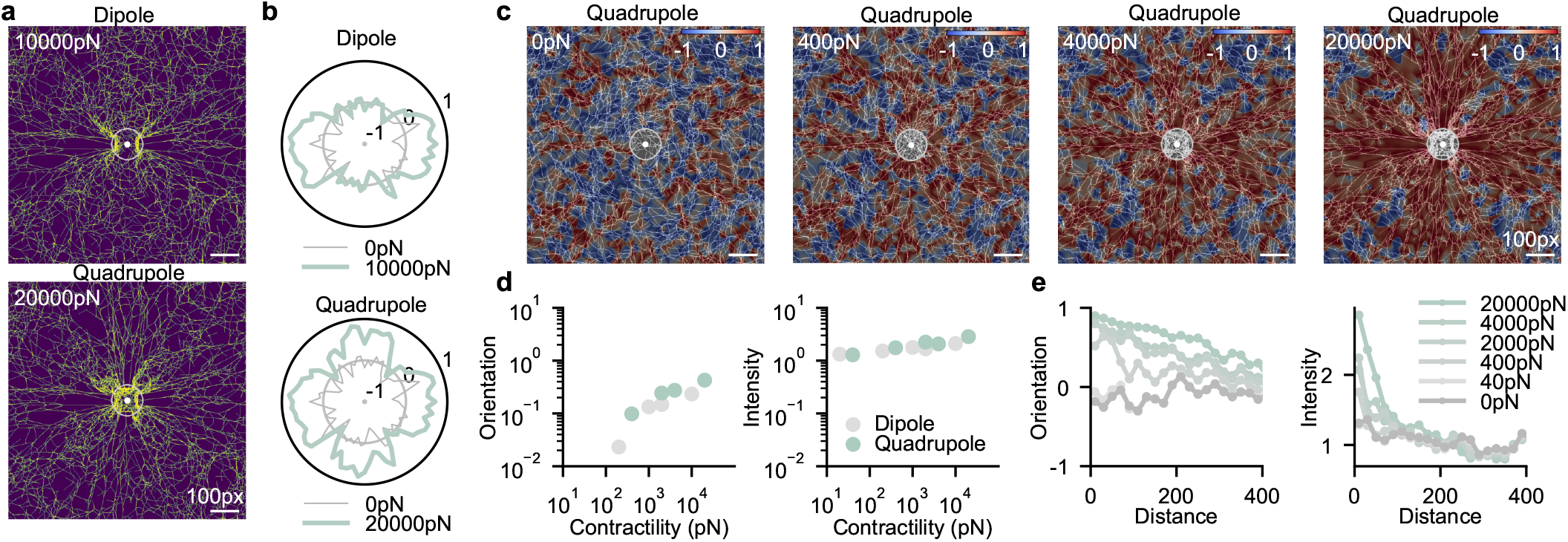
Fiber network model. **a**, Fiber network simulations (green) with contractile cells modelled as force dipoles (top, 2 point forces of 5000 pN, distance approx. 125 px) or quadrupoles (bottom, 4 point of 5000 pN, distance approx. 125 px). Contractility refers to the sum of individual point forces. **b**, Angular distribution of fiber orientation around a dipole or quadrupole (light green, cones bin width 5°), or for a cell-free network (gray). **c**, Fiber networks (black/white) and resulting orientation fields (blue/red) for a series of differently contractile quadrupoles. The mean fiber orientation (**d**, left) and fiber intensity close to the cell surface (**d**, right) increases with contractility similarly for both dipoles and quandrupoles. **e**) Fiber orientation (left) and fiber intensity (right) as a function of the distance from the outer cell contour, for dipoles (gray) and quadrupoles (green) and different contractilities.

We then apply forces of various magnitudes in a dipole or quadrupole arrangement to mimic contractile cells. As a result, fiber network densification and reorientation emerge that follow the shape of the force dipoles or quadrupoles (Fig. 4a-c). Increased force results in increased orientation and local densification (intensity) (Fig. 4d). Both the orientation and the intensity decrease with the distance from the cell, with orientation decaying more slowly (Fig. 4e). For a given force magnitude, the orientation and intensity values around dipoles and quadrupoles are similar, demonstrating that both scalar representations of force-induced fiber remodeling are largely insensitive to the details of how the forces are spatially arranged.

### Cell contractility correlates with fiber alignment and density

Next, we test if the positive, monotonic relationship between cell contractility and the surrounding collagen fiber orientation predicted by our simulated fiber network model as well as by previously published models (26, 27) can be experimentally verified. For this, we measure 3D traction forces (25) of three differently contractile primary glioblastoma cell lines embedded in collagen hydrogels (Fig. 5a, left) (36–39). The WK1 cell line (classical subtype) exhibit larger forces compared to the RN1 (mesenchymal subtype) and JK2 (proneural subtype) cell line (39). As 3D traction force measurements involve confocal imaging of the collagen matrix around each cell, we use the same data set to also evaluate the collagen fiber orientation intensity around the cell surface in distance shells spaced 10 μm apart. To compare measurements of cells obtained at different depths within the collagen gel, which can change the overall brightness of the image, the intensity within the first distance shell is normalized by the average intensity of the two outermost shells (Fig. 2a and SI Fig. 1).

**Fig. 5.**
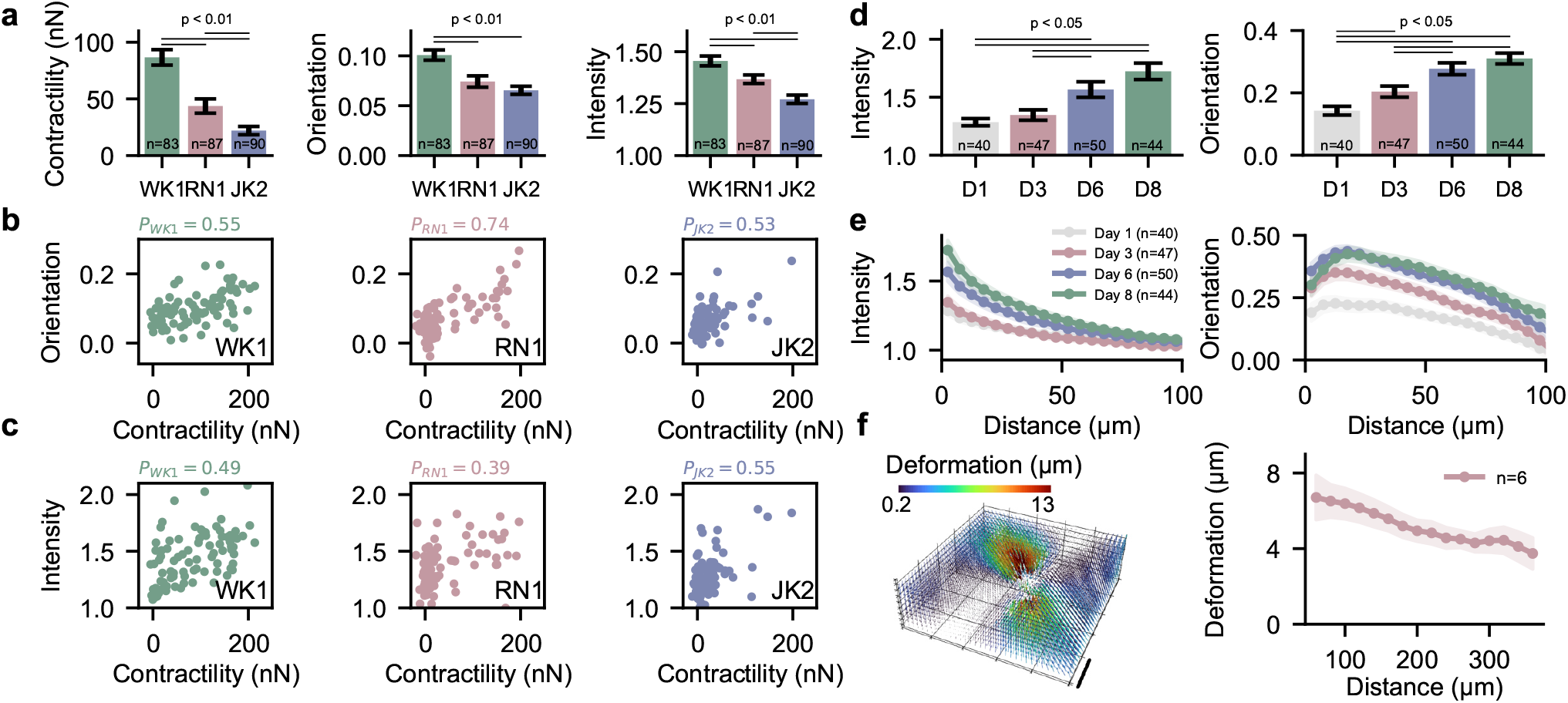
Cell contractility correlates with collagen fiber orientation and collagen intensity. **a**, Contractility, collagen orientation and collagen intensity (mean±se) for three different primary glioblastoma cell lines (WK1, RN1, JK1). n denotes the number of individual cells. **b**, Correlation of cell contractility with fiber orientation for three glioblastoma cell lines. P indicates Pearson correlation coefficient. **c**, Correlation of cell contractility with relative collagen fiber density (as measured from the local image intensity at a distance of 5 μm around the cell outline) for three glioblastoma cell lines. P indicates Pearson correlation coefficient. **d**, Time evolution of the relative increase in collagen fiber density (from the local image intensity at a distance of 5 μm around the cell outline; mean±se, left) and of the fiber orientation averaged over the image (mean±se, right) over a time course of 8 days after seeding hepatic stellate cells in collagen gels. **e**, Relative collagen fiber density (left) and fiber orientation (right), averaged over 5 μm wide areas around the cell outline, for different distances from the cell surface, for different days after seeding hepatic stellate cells in collagen gels. Data points represent the mean, shaded areas indicate mean±se. **f**, Matrix deformations around a representative hepatic stellate cell (left) embedded in 1.0 mg/ml collagen for 3 days. 95 percentile of the matrix deformations calculated over 20 μm wide shells as a function of distance from the deformation epicenter (right). Scalebar indicates 100 μm. Data points represent the mean, shaded areas indicate one standard error around the mean.

Both, collagen orientation and intensity increase with higher contractility (Fig. 5a). Additionally, we observe a statistically significant positive correlation between collagen orientation and contractility of individual cells for all three cell types (Pearson correlation coefficient P=0.61±0.09, mean±sd; Fig. 5b). We also observe a statistically significant positive correlation between collagen intensity and contractility (P=0.48+0.07, mean±sd; Fig. 5c). Although both findings qualitatively confirm the model predictions of a close relationship between contractility, fiber orientation and fiber intensity, the substantial variability in the data and the nonlinear shape of the relationships indicate that fiber orientation or intensity cannot quantitatively predict the contractility of an individual cell.

### Matrix remodelling over time

Next, we investigate the time course of matrix remodeling after cell seeding. The orientation and intensity of collagen fibers around HSCs are measured at 1, 3, 6, and 8 days after seeding. We observe that both quantities increase with culture time (Fig. 5d,e). Time-lapse imaging demonstrates that hepatic stellate cells progressively contract and align collagen over time (SI Video 16–19).

Contrary to model predictions, we observe that the fiber orientation radially increases over a distance of 10-15 μm from the cell surface and only begins to decrease at further distances (Fig. 5e, right). This discrepancy may be explained by local details of how cells interact with the surrounding matrix, as well as by fiber remodelling processes e.g. due to proteolytic processes or the secretion of matrix fibers. All of these processes are expected to modulate how the collagen structure locally reorients and remodels in response to local forces. The measured fiber intensities close to the cell surface, however, agree with the model predictions (Fig. 4e). Together, this suggests that for maximum robustness, fiber intensities should best be analyzed close to the cell surface, and fiber orientation should best be analyzed at distances larger than 20 μm away from the cell surface.

### Long-range propagation of matrix remodelling for drug-screening assays

In a non-linear fibrous matrix that deforms locally in a non-affine manner, such as collagen or fibrin, matrix deformations propagate over much further distances compared to linear elastic materials (25, 38, 40). To explore the relationship between cell contractility and long-range propagation of matrix deformations, fiber orientation, and fiber density (intensity), we measure these parameters for hepatic stellate cells cultured for 2 days in collagen gels. Fiber orientation and intensity are evaluated in intensity projected image stacks around the centered cells (Fig. 6e,h) as well as in 10 μm spaced distance shells up to a distance of 150 μm around the cells (Fig. 6f,i). Cells are cultured in the absence (con-trol) and presence of 10 μM Rho-kinase (ROCK) Inhibitor Y-27632. Rock-inhibition is known to reduce contractile force generation, and the Rock-pathway is of particular interest for liver fibrosis where it has been linked to the activation of hepatic stellate cells (41).

**Fig. 6.**
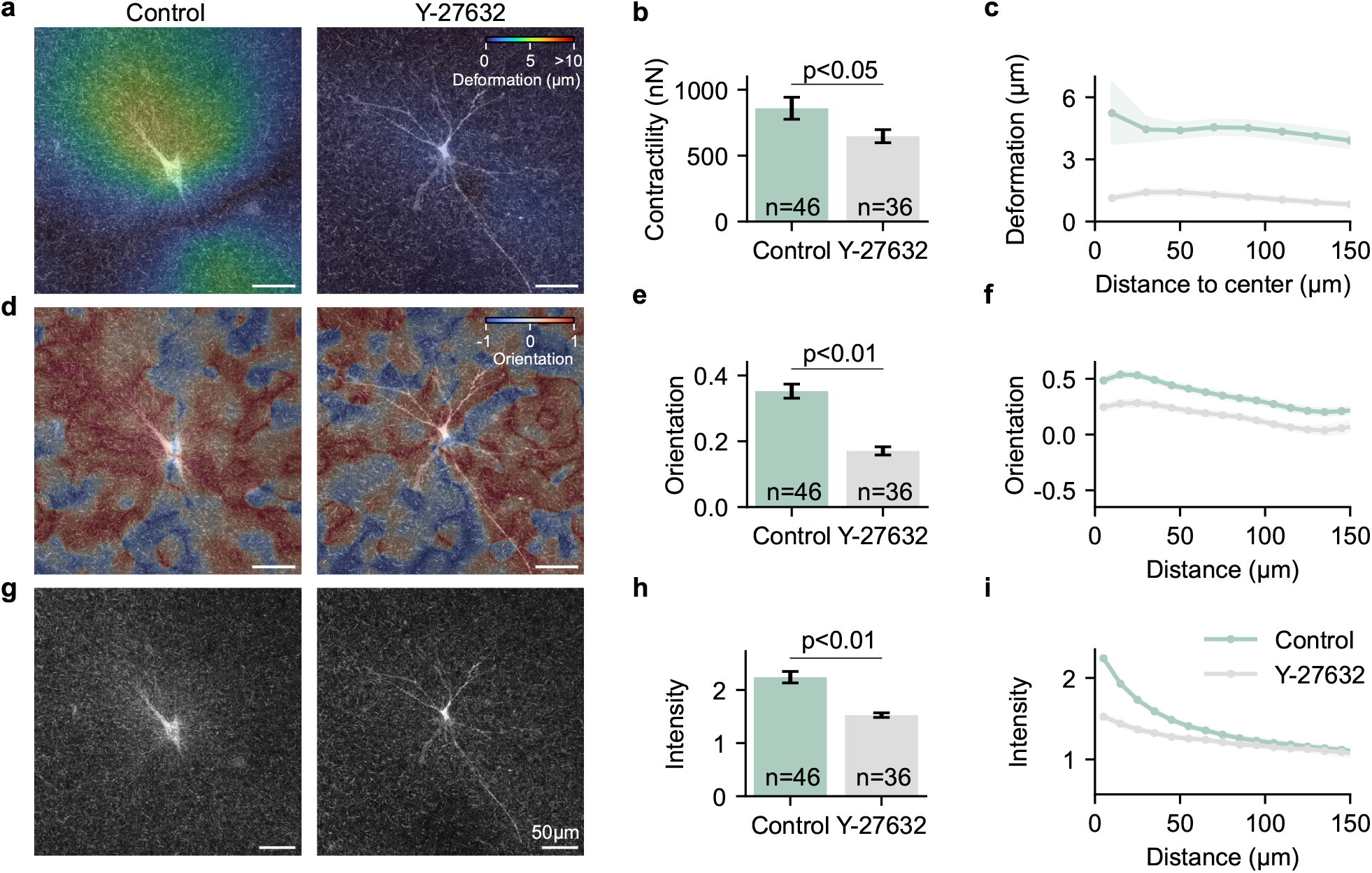
Matrix remodelling as a drug screening assay. Hepatic stellate cells are embedded in 1.2 mg/ml collagen hydrogels and cultured without (left) and with the addition of 10 μM Rock-Inhibitor Y-27632 (right) for 2 days. **a**, Images show an overlay of maximum intensity projections of the collagen (20 μm z-height of confocal reflection signal) around the cell and maximum intensity projections of the calcein stained cell. Colormap shows matrix deformations calculated using an optical flow algorithm from images taken before and after actin depolymerization with cytochalasin D (42) for a representative cell for each condition (see Suppl. Fig. 6 for more cells). **b**, From the 3D matrix deformations, the cell contractility is computed using traction force microscopy for control cells and cells treated with 10 μM Rho-kinase (Rock) Inhibitor (Y-27632). A significantly lower contractility is observed for cells treated with Rock-Inhibitor (mean±se; independent sample t-test). **c**, 3D Matrix deformations are averaged in 20 μm sized distance shells around the force center. **d**, Similar to **a**, color indicates the collagen fiber orientation of control cells and cells treated with 10 μM Rock-Inhibitor. **e**, Averaged collagen fiber orientation shows significantly lower values for cells treated with Rock-Inhibitor (mean±se; independent sample t-test). **f**, Fiber orientation is averaged in 10 μm sized distance shells around the cell center. **g** Similar to **a**, increased collagen intensity is observed for control cells. **h**, Averaged collagen intensity (intensity close to the cell surface (10 μm) normalized by the outer collagen intensity) shows a significantly lower value for cells treated with 10 μM Rock-Inhibitor (mean±se; independent sample t-test). **i**, Fiber intensity (normalized by the outer collagen intensity) is averaged in 10 μm sized distance shells around the cell center. Y-27632 treated cells with lower contractility, fiber orientation and fiber intensity also exhibit pronounced morphological differences compared to the more compact control cells (more cells shown in SI Fig. 6).

We find that matrix deformations propagate over long distances around hepatic stellate cells (Fig. 5f, 6c). Collagen fiber orientations (Fig. 5e, 6f) propagate in a similar manner, in agreement with the simulated fiber network model (Fig. 4e, green line: 20 nN), and also consistent with our findings in glioblastoma cells (SI Fig. 4,5). The modelled intensities (Fig. 4e) and the measured intensities (Fig. 5e, 6i) agree equally well; both decrease faster with distance compared to fiber orientation.

After two days of culture, we find that cells treated with 10 μM Rho-kinase (Rock) Inhibitor (Y-27632) show significantly lower contractile forces compared to control cells (Fig. 6b). In line with this finding, we observe that also the collagen fiber orientation and fiber intensity are significantly (p < 0.01) smaller (Fig. 6, SI Fig. 6). This demonstrates that both fiber orientation and fiber intensity are sensitive to drug-induced changes in cell contractility.

Since measurements of 3D traction forces require the relaxation of cell forces by adding the actin depolymerizing agent cytochalasin D (SI Video 15), we can quantify the residual fiber orientation and intensity in the matrix under force-free conditions. After drug-relaxation, the fiber orientation and intensity relax by approx. 30% (SI Fig. 7). This indicates the presence of strong remodelling processes already after 2 days of culture, e.g. due to proteolytic activities, plastic deformations, or crosslinking. This is of particular interest for comparing fiber orientation and intensity measurements with traction force microscopy: traction force methods account only for elastically stored forces in the matrix that are released within a short time period after addition of a force-relaxing drug. Thus, traction force methods ignore the forces that may have induced non-elastic deformations in the matrix over a prolonged culture time. By contrast, fiber orientation and intensity measurements report both, elastic and non-elastic matrix deformations due to cell-generated forces. Estimation of cell-generated forces from the local fiber orientation and and fiber density around cells is not limited to collagen hydrogels, but can also be applied to other fibrous matrices, such as fibrin gels. We observe a similar propagation of fiber orientation and fiber density (SI Fig. 8) for human lung fibroblasts cultured in fibrin gels.

In summary, fiber orientation and fiber intensity around contractile cells can be used for qualitatively estimating cellular force generation in fibrous 3D matrices. The method is particularly suitable for drug screening assays where the absolute values of cell contractility are less relevant and only drug-induced differences in cell contractility are of interest. In contrast to 3D TFM, fiber orientation and fiber intensity measurements are relatively simple. They do not require the addition of a force-relaxing compound, which allows for higher throughput and repeated measurements over time of the same cells or on the same sample. To analyze the fiber images and extract the relevant information, we have developed an open source Python package (29) with a graphical user interface (SI Fig. 9).

## Discussion

In this study, we quantify matrix remodeling around contractile cells in 3D fiber matrices by measuring the fiber orientation and fiber density. Using fiber network models and cell experiments in collagen fiber networks, we demonstrate both theoretically and experimentally a correlation between fiber orientation, fiber intensity, and cell contractility. Exploiting this correlation allows us to interpret fiber remodelling as a proxy for cell contractility. We demonstrate that the method can be applied as a drug-screening assay to qualitatively estimate changes in cell contractility. This method does not require knowledge of the matrix mechanical properties, and is experimentally and computationally simpler compared to classical traction force measurements.

Fiber orientation and intensity are both sensitive to changes in cell contractility, but to a different degree. Collagen and fibrin fiber orientation around contracting cells shows long-range propagation, similar to fiber deformation, whereas fiber intensity decreases more rapidly with distance. This implies that when analyzing the far field, fiber orientation - although it is more difficult to quantify than intensity - may provide a more robust measure of cell contractility. However, close to the cell, fiber intensity is more sensitive to cell contractility compared to fiber orientation, which shows a non-monotonic behavior in the form of an initial increase over a ∽20 μm distance before it decreases (Fig. 5, 6, SI. Fig. 4). This behavior arises because we compute the fiber orientation relative to the cell center, which averages the contribution of the individual cell protrusions. In addition, accumulated or secreted collagen near the cell surface may also interfere with the orientation analysis.

While a sufficiently large size of the imaged matrix in the horizontal (x,y) plane is important for resolving the far-field, the height (z-direction) of the imaged volume around the cell needs to be sufficiently small. For our analysis, we use maximum intensity projections to incorporate collagen fibers from different layers around individual cells, with a total z-height between 10-50 μm, depending on the cell type. If the z-height of the imaged volume is too large, undeformed collagen outside the cell plane may reduce the overall sensitivity, while for a height that is too small, regions of re-modeled collagen may be missed.

For different experimental setups and matrices (collagen, fibrin), a suitable window size for the analysis must be chosen. A good starting point for the window size is the pore size of the fiber network. This window size can then be optimized by finding the size at which the orientation value becomes a maximum. A relationship between orientation and window size for a typical example image is shown in (SI Fig. 2). The Python implementation of our method provides an algorithm for automatically determining the optimal window size.

In addition to estimating contractile forces, quantification of altered matrix remodeling may also be of interest for studying disease processes such as fibrosis and cancer (13–15, 43–47). For example, so-called Tumor-Associated Collagen Signatures (TACS) have been introduced as prognostic markers in cancer. The metric TACS-1 denotes an increased accumulation of collagen around the tumor, which is related to the fiber intensity in our approach. TACS-2 denotes stretched collagen in proximity to the tumor surface as the tumor grows, which is related to the fiber orientation in our approach (13, 43–46, 48). In fibrosis, cells transmit mechanical signals over long distances, and mechanically mediated cell-cell communication plays a role in the activation of quiescent cells (49–54). In particular the fiber orientation, which is sensitive to matrix remodelling in the far-field, may therefore be a useful metric to explore and quantify fibrotic disease progression and cell-cell-communication (see SI. Fig.10,11, SI Video 17,19 for breast cancer and liver fibroblasts).

Compared to 3D traction force microscopy, our method allows for higher throughput, since samples do not need to be relaxed by additional drug treatment (e.g. cytochalasin D), and is also suitable for repeated measurements of the same sample over long periods of time or for multicellular spheroids (SI Fig. 11). The method is available as an open-source Python package (29) together with a graphical user interface (SI Fig. 9).

## Material and Methods

### Cell Culture

#### Hepatic stellate cells (HSC)

Primary human cryopreserved hepatic stellate cells (HUCLS, Lonza, Basel) are cultured at 37°C, 95% humidity and 5% CO_2_ in human hepatic stellate cell growth medium (MCST250, Lonza, Basel).

#### Human Lung Fibroblasts (NHLF)

Normal Human Lung Fi-broblasts (NHLF, Lonza, Basel) are cultured at 37°C, 95% humidity and 5% CO_2_ in Fibroblast Growth Medium (FGM-2, Lonza, Basel).

#### Glioma neural stem cells (GNS)

Patient-derived glioma neural stem cells (WK1, RN1, JK2) are provided by the QIMR Berghofer Medical Research Institute in Brisbane (36, 37, 55). Cells are cultured at 37°C, 95% humidity and 5% CO_2_ in knockout DMEM/F-12 medium supplemented with recombinant human EGF (20 ng/ml), recombinant human FGFb (10 ng/ml), glutamine (20 mM/ml), penicillin/streptomycin (100 U/ml) and StemPro neural supplement (20 ng/ml), all from Thermo Fisher Scientific, Waltham, as well as heparin (20 ng/ml, Sigma-Aldrich, St. Louis). Cell culture flasks are coated with 1% growth factor-reduced Matrigel (BD Biosciences, Mississauga) diluted in DMEM medium (high glucose, Thermo Fisher Scientific, Waltham).

### Fiber orientation and intensity analysis

The fiber orientation and intensity analysis are based on a maximum intensity projected image of the fiber network, and a maximum intensity projected image of the cell, computed from image stacks with variable size (specified below for the individual experiments). Cell area and position are segmented from the cell image using contrast enhancement, Gaussian filtering, morphological operations and Otsu’s thresholding (56).

The cell area is excluded from further analysis. The fiber image is contrast-enhanced and Gaussian-filtered (sigma = 0.5 pixel). The predominant matrix fiber orientation is determined based on structure tensor analysis (30–32), implemented similar to previously described approaches (33–35). Specifically, the local structure tensor *S*_0_ is computed from the partial derivatives of the pixel intensities in the x-direction (*f*_*x*_) and y-direction (*f*_*y*_) of the image according to Eq. 1 (30–32). At each pixel *p*, we use the local structure tensor *S*_0_ in a surrounding region by convolution with a Gaussian kernel *w*. Structural features with much smaller length-scales compared to the size of this region are therefore not contributing to the structure tensor *S* (Eq. 2). The size of the convolution region is defined by the standard deviation σ (also referred to as window-size) of the Gaussian kernel *w* (Eq. 3). In this study, the window-size is set in the range of 7-10 μm, which is about twice the collagen pore size of 5.5 μm (SI Fig. 2, values specified below for individual experiments).

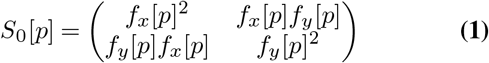

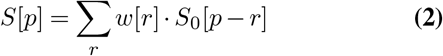

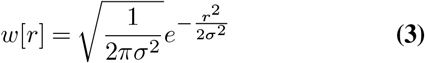

From the structure tensor *S*, we compute the corresponding eigenvectors 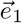 and 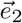 that encode the orientation vector of the structures (e.g., of a matrix fiber). The eigenvector belonging to the larger eigenvalue indicates the direction with the highest pixel intensity gradient of the structure, whereas the eigenvector belonging to the smaller eigenvalue points perpendicular to the gradient direction and therefore is the orientation vector of the structure 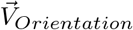 (Eq. 4) (57).

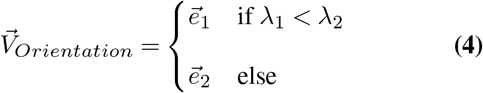

Based on the eigenvalues *λ*_1_, *λ*_2_, we can also quantify the anisotropy of the structure orientation, also referred to as coherence *c*_*w*_ (Eq. 5) (32, 33).

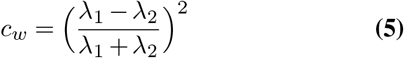

A coherency of 0 indicates isotropically oriented structures, whereas a value of 1 indicates perfect alignment (32). Next, we calculate the angular deviation between the structure orientation *V*_*Orientation*_ and the vector pointing towards the cell center *V*_*Center*_ from the magnitude of the scalar product between the two vectors. The angular deviation is then transformed into a orientation value Θ_*i*_ that ranges from -1, if the structure is perpendicular to the center-line, up to 1, if the structure is parallel to the center-line. A value 0 corre-sponds to a orientation of 45° (Eq. 6).

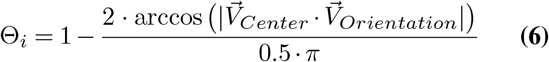

If the the local intensity or the local coherency of the fiber image is low, the local orientations Θ_*i*_ cannot be computed with high confidence. Therefore, when averaging Θ_*i*_ for each cell, the local Θ_*i*_ values are weighted by the local coherence *c*_*w,i*_ and the local image intensity *I*_*w,i*_ according to Eq. 7. In addition, regions near the image borders (40 px) and the cell-occupied area are excluded from the analysis.

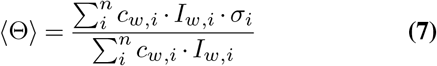

Local collagen orientation values around each cell are then averaged over the entire image (Fig. 2c,d, SI Fig. 3), averaged over areas bounded by equidistant lines (“shells”) around the cell contour (Fig. 2a), or averaged over azimuthal bins (“cones”) around the cell center (Fig. 4b, SI Fig. 1). Further, the collagen intensity is normalized to the mean collagen intensity within the two most distant shells. The normalized intensity in the first distance shell, close to the cell surface, is then used as measure of the increased fiber intensity per cell.

### Artificial line networks

Spherical artificial cells (diameter 200 px, n = 30) are placed in images (1024×1024 px) with diagonal, radial and concentric lines around the cell center. The line widths in the images are varied between 1 px and 5 px (step size: 1 px). The number of fibers is varied between 50 and 100 (step size: 10). Additionally, artificial cells are placed in images containing Gaussian noise (n = 30). The orientation analysis is performed using a window-size of 10 px.

### Simulated fiber networks

The fiber network simulations are set up as described in our previous studies (58, 59). Briefly, fibers are polymerized from monomeric units (800 nm long cylindrical segments) in random directions during network formation. Neighboring fibers are interconnected by crosslinks. Fibers segments have a bending stiffness (spring constant) of 7.54·10^−16^ Nm^-1^ and an extensional stiffness (spring constant) of 0.055 Nm^-1^. Cell tractions are simulated by either a dipole or a quadrupole placed in the center of the simulation domain with specified force magnitudes. The simulation domain size is 200×200×1 μm for computational feasibility. The distance between point forces is approx. 125 px (30 μm). Fiber orientation analysis is performed using a window-size of 10 px (approx. 2.4 μm). Polar averaging is performed using 5°-sized angle sections (“cones”). The fiber orientation and intensity propagation are evaluated in 20 px (approx. 4.8 μm) spaced distance shells around the cell. The intensity is normalized to the intensity of the two outermost shells within the field of view.

### Glioma neural stem cells (GNS) experiments

#### Single cell traction force analysis

Traction force experiments of glioma neural stem cells are conducted as described previously (38). In brief, collagen type I hydrogels are prepared using a 1:2 mass ratio of acid-dissolved rat tail (R) and bovine skin (G1) collagen (Matrix Bioscience, Mörlenbach) dissolved in a solution of 1 vol part NaHCO_3_, 1 vol part 10×DMEM and 8 vol parts H_2_O. The pH of the solution is adjusted to 9 by adding NaOH (1 M). Cells and collagen solution are transferred to a 35 mm dish (bottom layer: 1.75 ml collagen pre-polymerized for 2.5 minutes, top layer: 250 μl collagen containing 15.0000 cells) and polymerized at 37° for 60 minutes at a final collagen concentration of 1.2 mg/ml. Subsequently, 2 ml of cell culture medium are added, and after an additional waiting time of at least two hours, a cubic volume V=(370 μm)^3^ (voxel-size 0.72× 0.72×0.74 μm) around individual cells is imaged in confocal reflection and brightfield mode using an upright confocal laser scanning microscope equipped with a 1.0 NA 20x dip-in objective (Leica SP5, Leica, Wetzlar). The relaxed reference state of the collagen is acquired 30 minutes after cytochalasin D treatment (20 μM). Cell-generated forces are calculated from the resulting matrix deformations as described in (38) using the elastic network optimizer Saeno (25). Three independent experiments are conducted for each cell type.

#### Analysis of matrix remodelling

Maximum intensity projected images of the collagen fiber network (confocal reflection mode) and the cell outline (brightfield mode) are calculated using 20 μm (z-height) stacks around the cells. Cells are segmented based on the projected brightfield image using Otsu-thresholding (60), and the fiber orientation is calculated from the projected confocal reflection images as described above. The relative increase in collagen fiber intensity around the cell is calculated from the first 10 μm distance shell normalized to the two outermost shells. For the collagen orientation analysis, a window-size of 10 μm is chosen.

### Hepatic stellate cells (HSCs) experiments

#### Analysis of matrix remodelling (time course analysis, Fig.5d,e)

Collagen type I hydrogels (RAFT Reagent Kit for 3D culture, Lonza, Basel) are prepared on ice to a final collagen concentration of 1.0 mg/ml. The collagen solution is mixed with hepatic stellate cells and transferred to a 96-well plate (μ-Plate, Ibidi, Gräfelfing), seeding 4000 HSCs in 200 μl collagen solution per well. Gels are polymerized at room temperature (RT) for 10 minutes, after which 350 μl of HSC-medium is added per well. Prior to imaging, cells are stained with calcein (Celltrace Calcein Red-Orange AM, Invitrogen, Waltham). 100 μl medium is removed from each well and replaced with 100 μl calcein-medium solution for a final calcein concentration of 1.5 μM. Samples are incubated for 15 min at 37°C, gently washed 3 times with medium (leaving a 100 μl medium layer over the gel surface) and covered with 250 μl of fresh medium. Samples are transferred to a multiphoton excitation microscope equipped with a 1.0 NA 25x objective (FVMPE-RS, Olympus, Tokyo). Image stacks (xy= 255×255 μm, 0.318 μm per pixel) are recorded around individual cells using bright-field mode, second-harmonic generation, and calcein fluorescence. For experiments shown in Fig. 5b, a stack height of 70 μm with a z-step of 2 μm is recorded. Cells are segmented based on the projected calcein-stained image, and the fiber orientation is calculated based on the projected confocal reflection image of the surrounding collagen. The collagen orientation and fiber intensities are calculated over 5 μm distance shells and normalized to the intensity of the two outermost shells within the field of view. A window-size of 7 μm is used for the collagen orientation analysis.

#### Particle image velocimetry (Fig. 5f)

Image stacks around individual cells (xyz = 509 × 509 × 148 μm, with a voxel-size of 0.636 × 0.636 × 2 μm) are recorded once before and 4 h after actin-depolymerization using 10 μM cytochalasin D (Sigma-Aldrich, St. Louis) using brightfield and second-harmonic generation imaging microscopy. 3D matrix deformations are then calculated between the contracted and the relaxed state of the collagen matrix using 3D Particle Image Velocimetry (61, 62). Here, a window-size of 35 μm together with 60% overlap between neighbouring windows (SI Fig. 12) and a signal-to-noise threshold of 1.3 are used. Deformation fields are drift corrected, and the deformation epicenter is determined as described in (25, 62). In order to minimize bias from noise, we only consider deformations that point ±60° towards the deformation epicenter. We calculate the 95 percentile of absolute matrix deformations over 20 μm distance shells around the cell center. For the data shown in Fig.6c, the force epicenter is used instead of the deformation epicenter.

#### Single cell traction force analysis (TFM, Fig. 6)

Traction force experiments of hepatic cells are conducted as described in (38, 62). Collagen type I hydrogels are prepared using a 1:2 mass ratio of acid-dissolved rat tail (R) and bovine skin (G1) collagen (Matrix Bioscience, Mörlenbach) that are dissolved in a solution of 1 vol part NaHCO_3_, 1 vol part 10 × DMEM and 8 vol parts H_2_O. The pH of the solution is adjusted to a value of 9 by adding NaOH (1 M). 35.000 cells are mixed with 3 ml collagen solution (final collagen concentration of 1.2 mg/ml) and transferred to a 35 mm Petri dish (Greiner AG, Austria) and polymerized at 37° for 60 minutes. Subsequently, 2 ml of cell culture medium containing either 10 μM Rho-kinase (Rock) Inhibitor Y-27632 (Sigma-Aldrich, St. Louis) or identical amounts of PBS (control) are added. Two days after embedding, the medium is replaced (with fresh Rock-Inhibitor or PBS). Prior to imaging, cells are stained with 2 μM calcein AM (Thermo Fisher Scientific, Waltham) and incubated for 20 min. Samples from three independent experiments are imaged with an upright confocal laser scanning microscope (Leica SP5, Leica, Wetzlar) equipped with a 20x dip-in objective (NA 1.0, Leica, Wetzlar) at 37°C and 5% CO_2_. A cubic volume V=(370 μm)^3^ is imaged around individual cells in confocal reflectance, brightfield, and fluorescence modes (voxel-size 0.72×072×0.99 μm/pixel). The force-free reference state of the collagen is recorded 30 minutes after actin-depolymerization with 10 μM cytochalasin D (Sigma-Aldrich, St. Louis). 3D matrix deformations are quantified using Particle Image Velocimetry (OpenPIV) with a window-size of 35 μm (SI Fig.12) and an overlap of 60% between neighbouring windows (61, 62). From the drift-corrected deformation fields, we compute the cell-generated forces using the software Saenopy (62), a Python implementation of the elastic network optimizer Saeno (25). Force regularization is performed using a regularization parameter of *α* = 10^10^, with nonlinear elastic material parameters for 1.2 mg/ml collagen gels (linear stiffness *k*_0_=6062, linear strain range *λ*_*s*_=0.0804, stiffening coefficient *d*_*s*_=0.034, and buckling coefficient *d*_0_=0.0025 (62)).

#### Analysis of matrix remodelling (TFM Data, Fig. 6)

Max-imum intensity-projected images of the collagen fiber networks (confocal reflection mode) and the calcein-stained cells (confocal fluorescence) are calculated from 20 μm (z-height) image stacks. Cells are segmented using Otsu-thresholding (60), and the fiber orientation is calculated from the projected confocal reflection images as described above. Relative increase in collagen fiber intensity around the cell is calculated from the first 10 μm distance shell normalized to the two outermost shells. For the collagen orientation analysis, a window-size of 10 μm is chosen.

### Lung fibroblasts (NHLF) experiments

Normal Human Lung Fibroblasts (NHLF, Lonza, Basel) are embedded in fluorescently labeled fibrin gels as previously described (58). NHLF are labeled with CellTracker (30 min incubation at a concentration of 0.66 μM, CellTracker Green CMFDA, Thermo Fisher Scientific, Waltham). Bovine fibrinogen is labeled with a fluorescent reactive dye (Alexa Fluor 647 NHS succinimidyl ester, Thermo Fisher Scientific, Waltham). After purification from the unreacted dye via dialysis, the fibrinogen (6 mg/ml) is mixed 1:1 with bovine thrombin (2 unit/ml) and added to a NHLF cell pellet for a final concentration of 50,000 cells/ml, 3 mg/ml fibrinogen and 1 unit/ml thrombin. The mixture of fibrin gels and NHLFs is pipetted into PDMS chambers (1.3 mm ×1.3 mm×130 μm) as previously described (58). After 4 hours of incubation, samples are transferred to a confocal microscope (SPE, Leica, Wetzlar) equipped with a 10x 0.4 NA objective (Leica, Wetzlar).

Confocal image stacks (xyz=256×256×58 μm, with a voxel-size of 0.5×0.5×0.5 μm/px) are acquired around individual cells. Maximum intensity projected images are calculated from 20 μm (z-height) stacks around the cells. Cells are segmented using Otsu-thresholding (60), and the fiber orientation is calculated from the projected confocal reflection images as described above. Relative increase in collagen fiber intensity around the cell is calculated from the first 10 μm distance shell normalized to the two outermost shells. For the collagen orientation analysis, a window-size of 10 μm is chosen.

### Spheroid experiments (CAFs)

Luminal B breast cancer associated fibroblasts (CAFs) are isolated from patients as previously described (38). Human tissue collection was approved by the Ethics Committee of the Friedrich-Alexander University Erlangen-Nürnberg, Germany (#99_15Bc) in accordance with the World Medical Association Declaration of Helsinki. Cells are cultivated in Epicult basal medium with Supplement C (Epicult-C human media kit, Stem Cell Technologies, Vancouver), 5% FCS, 1% penicillin, and 1% streptomycin. Multicellular spheroids are formed from 4000 cells in 100 μl medium by culturing for 1 day in non-adhesive U-well plates (Greiner, Kremsmünster). The next day, 8-12 spheroids are embedded in 1.2 mg/ml collagen gels (Matrix Bioscience, Mörlenbach) using a previously described two-layer gel approach (38) to prevent the spheroids from sinking to bottom of the dish (1.5 ml collagen solution per layer, first layer polymerized for 20 minutes).

### Characterization of collagen gels

Collagen type I hydrogels (RAFT Reagent Kit for 3D culture, Lonza, Basel) are prepared on ice to a final collagen concentrations of 1.0 mg/ml and polymerized at room temperature in a 96-well plate (200 μl volume per well, μ-Plate, Ibidi, Gräfelfing). Samples are transferred to a multiphoton excitation microscope (Olympus FVMPE-RS Microscope, Tokyo) equipped with a 25x objective (Olympus XLPLN25XSVMP, NA 1.0, Tokyo) and imaged at 37°C, 95% humidity and 5% CO_2_. Image stacks (xyz=255×255×50 μm, with a voxel-size of 0.249×0.249×0.5 μm) are acquired using second-harmonic imaging microscopy. Image stacks are binarized, and the 3D pore size is extracted as described in (63, 64).

## Data Availability

The software is available as an open-source Python package with a graphical user interface on GitHuB (29). Figures are created using the Python package Pylustrator (65) and MetBrewer (66). The data of this study are available upon request from the corresponding author.

## Author contribution

AB, DB, MR, EG and BF developed the analysis method. AB, RG and DB developed the fiber analysis software. MM performed the fiber network simulations. IM, LB and DB and performed HSC single cell experiments. AM conducted fibrin experiments. CM, GO and TG conducted and analyzed the glioblastoma traction force experiments. SB and DB conducted the rheological measurements. RS and PS established the primary CAF cell lines. DB conducted spheroid experiments. DB conducted the data analysis. DB and DK created figures and tables. BF and DB wrote the manuscript.

## ACKNOWLEDGEMENTS

This work was funded by grants from the German Research Foundation (DFG TRR-SFB 225 – project 326998133 – subprojects A01, B09, C02 and C05; STR 923/6-1), and the Emerging Fields Initiative of the University of Erlangen–Nuremberg. We thank the ENB Biological Physics program of the University Bayreuth for support, and Ricardo Henriques for the use of the latex layout.

---

**Supplementary Information 1:**
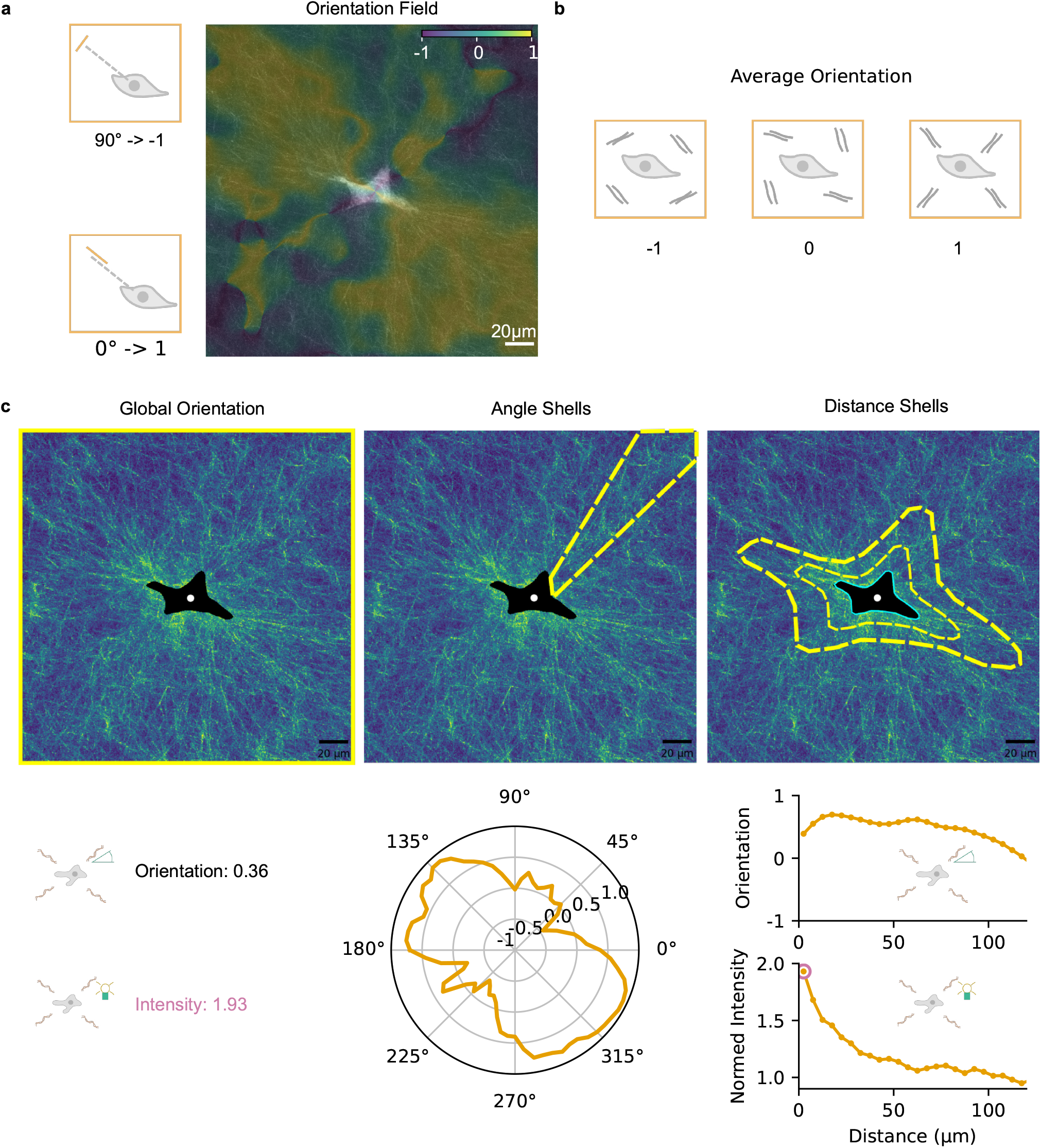
Analysis of matrix remodelling. **a**, The orientation of collagen fibers is quantified by computing the structure tensor from the maximum-projected fiber images (see Methods and Fig. 2). The fiber orientation with respect to the cell center is calculated according to Eq. 6. An orientation of -1 corresponds to fibers oriented perpendicular to the vector pointing to the cell center (90° deviation), and an orientation of +1 corresponds to elements pointing to the cell center (0° deviation). **b**,**c**, All orientation values are averaged within the field of view to obtain a global orientation value (**c** left), averaged within specified angular sections (cones, c middle), or averaged within distance shells around the cell contour (**c** right). The fiber intensity is normalized to the intensity of the two outermost distance shells. The normalized fiber intensity in the first (innermost) distance shell is used as a measure for the increased fiber density (**c** right).

**Supplementary Information 2:**
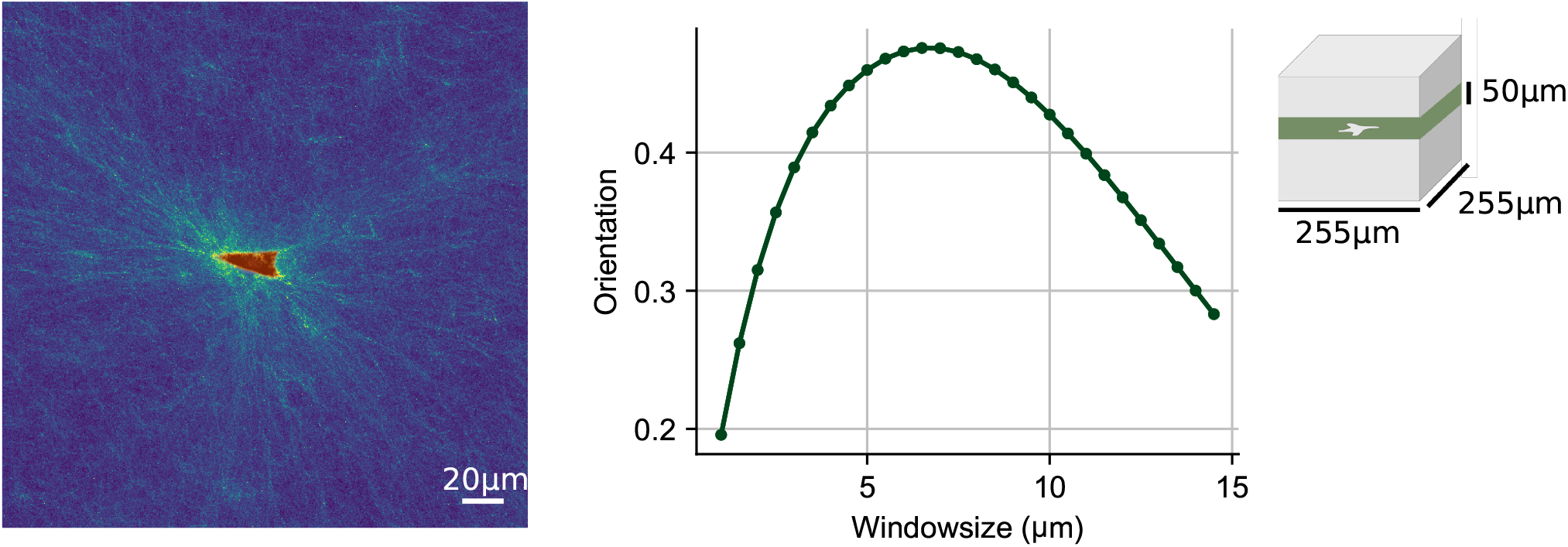
Determing a suitable window-size for the collagen orientation analysis. **Left**, Hepatic stellate cell cultured in 1.0 mg/ml collagen gels (Lonza, Basel) for 6 days. Image stack (xyz=255×255×50 μm) of the collagen is acquired by second harmonic generation microscopy (green-blue), and image stack of the calcein-stained cell is acquired using fluorescence microscopy (red). **Right**, Global collagen orientation is calculated from the maximum intensity projected fiber images using different window-sizes (see Methods section). Here, a window-size of 7 μm is chosen, which maximizes the collagen orientation and is comparable to the collagen pore size (5.5 μm).

**Supplementary Information 3:**
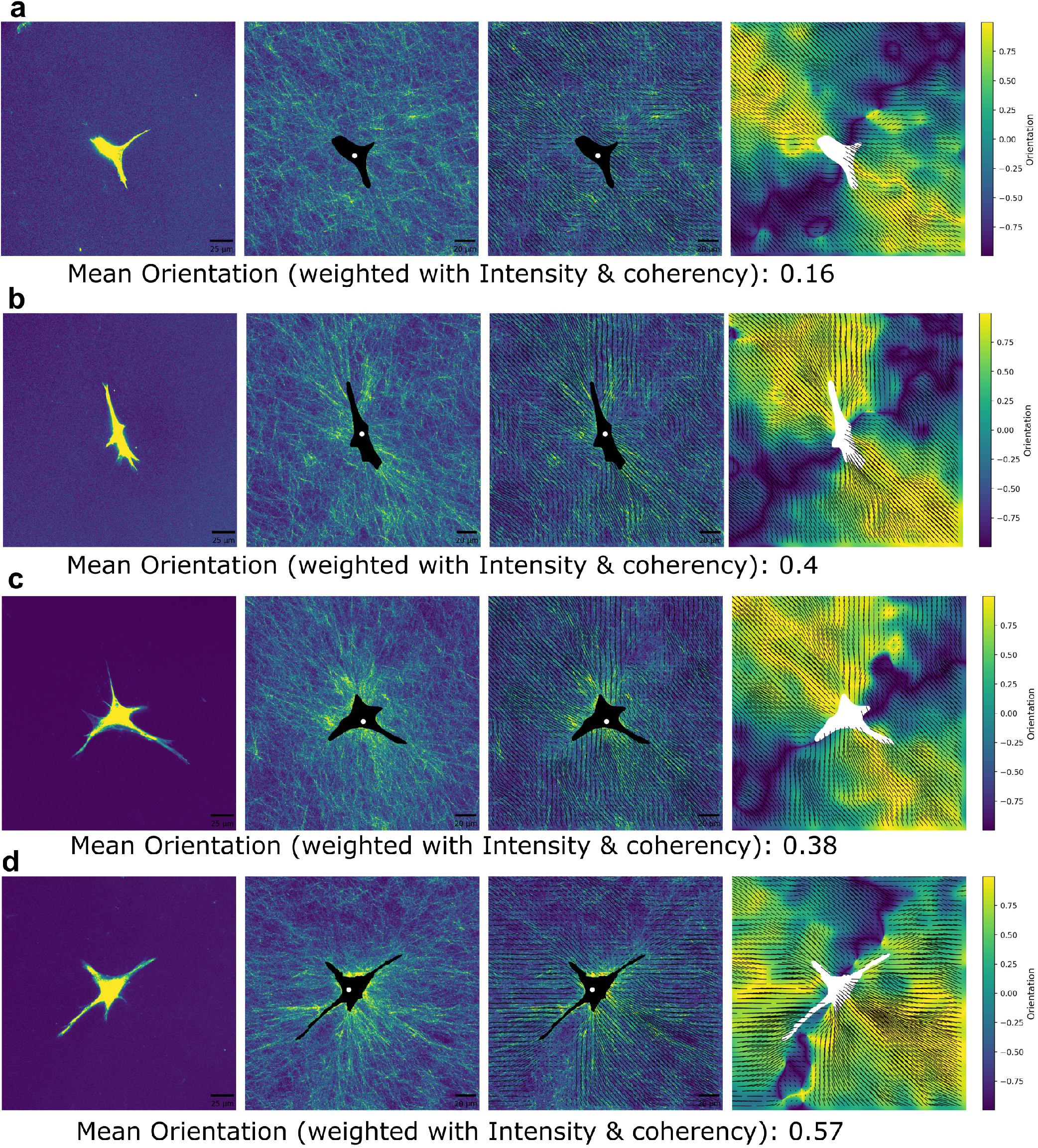
Collagen orientation around individual hepatic stellate cells. Four represantative hepatic stellate cells (**a-c**) imaged after 4 days of culture in 1.0 mg/ml collagen gels (**RT**, Lonza). First column: maximum intensity projected images of calcein-stained hepatic stellate cells. Second column: collagen fiber network measured using second harmonic generation imaging. The segmented cell area is shown in black, and the cell center is indicated by a white dot. Third column: orientation vectors (black arrows). Fourth column: orientation values with respect to the cell center according to Eq 6. Values below the images specify the averaged fiber orientation weighted by image intensity and fiber coherency.

**Supplementary Information 4:**
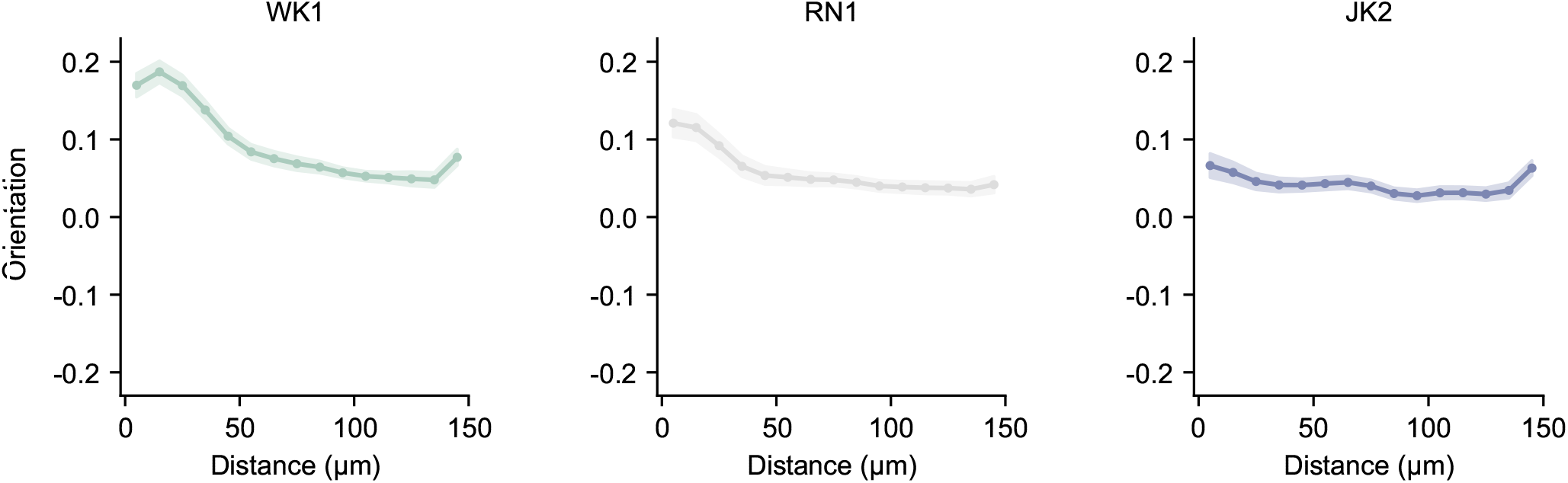
Fiber orientation around glioblastoma cells. Collagen fiber orientation around three different primary glioblastoma cell lines (WK1, RN1, JK1) in 1.2 mg/ml collagen gels calculated in 10 μm spaced distance shells around individual cells. Curves and shaded areas indicate mean±se. (Number of cells: WK1=83, RN1= 87, JK2= 90).

**Supplementary Information 5:**
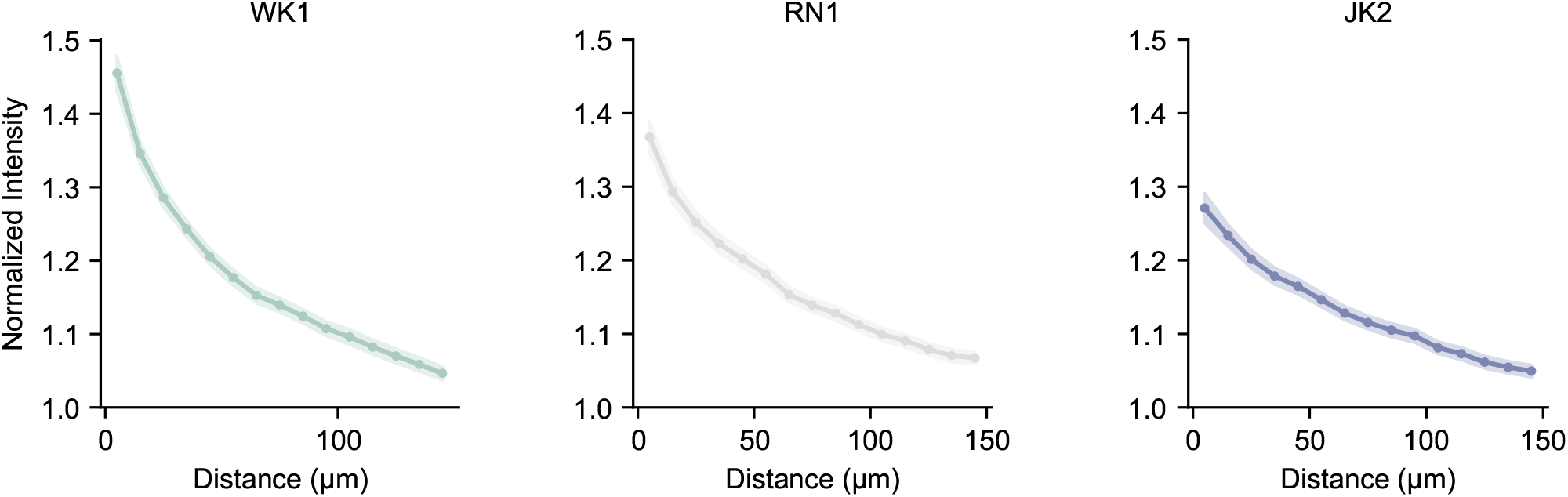
Fiber intensity around glioblastoma cells. Normalized collagen fiber intensity around three different primary glioblastoma cell lines (WK1, RN1, JK1) in 1.2 mg/ml collagen gels averaged within 10 μm distance shells around individual cells. Curves and shaded areas indicate mean±se. (Number of cells: WK1=83, RN1= 87, JK2= 90).

**Supplementary Information 6:**
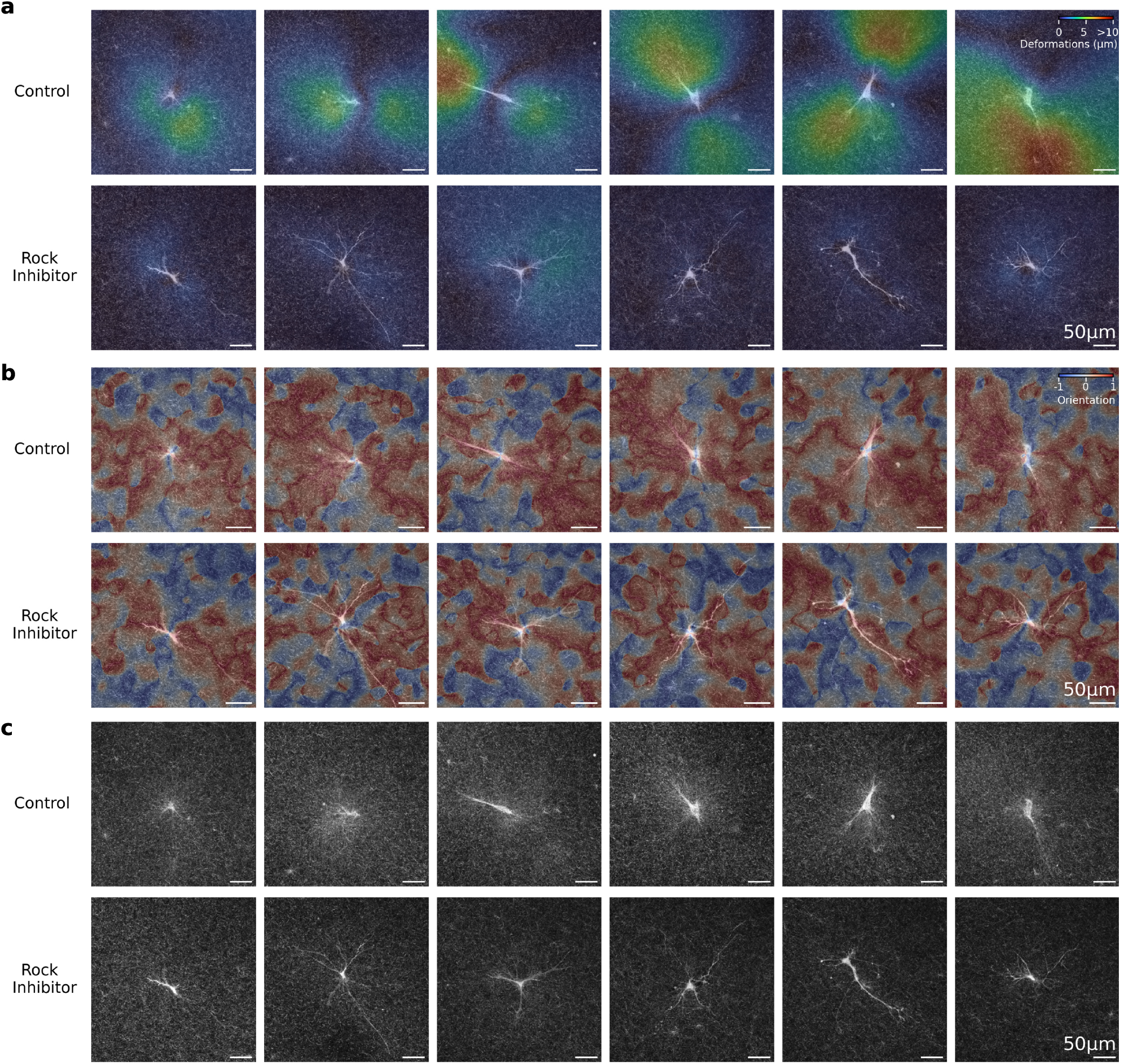
Matrix deformations around hepatic stellate cells in response to Rock-inhibition. Hepatic stellate cells are embedded in 1.2 mg/ml collagen gels (Matrix Bioscience, Mörlenbach) and cultured without (top) and with the addition of 10 μM Rho-kinase (Rock) Inhibitor Y-27632 (bottom) for 2 days. Images show an overlay of the maximum intensity projected collagen network (20 μm z-height around the centered cell) imaged with confocal reflection microscopy, and the maximum intensity projected calcein stained cell imaged with confocal fluorescence microscopy. **a**, Colormap indicates the matrix deformations between the projected collagen stacks before and after cytochalasin D treatment for six representative cells of each condition, calculated using an optical flow algorithm (42). **b**, Colormap indicates the collagen fiber orientation before cytochalasin D treatment for the same cells of each condition. **c**, Collagen fiber intensity is shown for the same cells of each condition.

**Supplementary Information 7:**
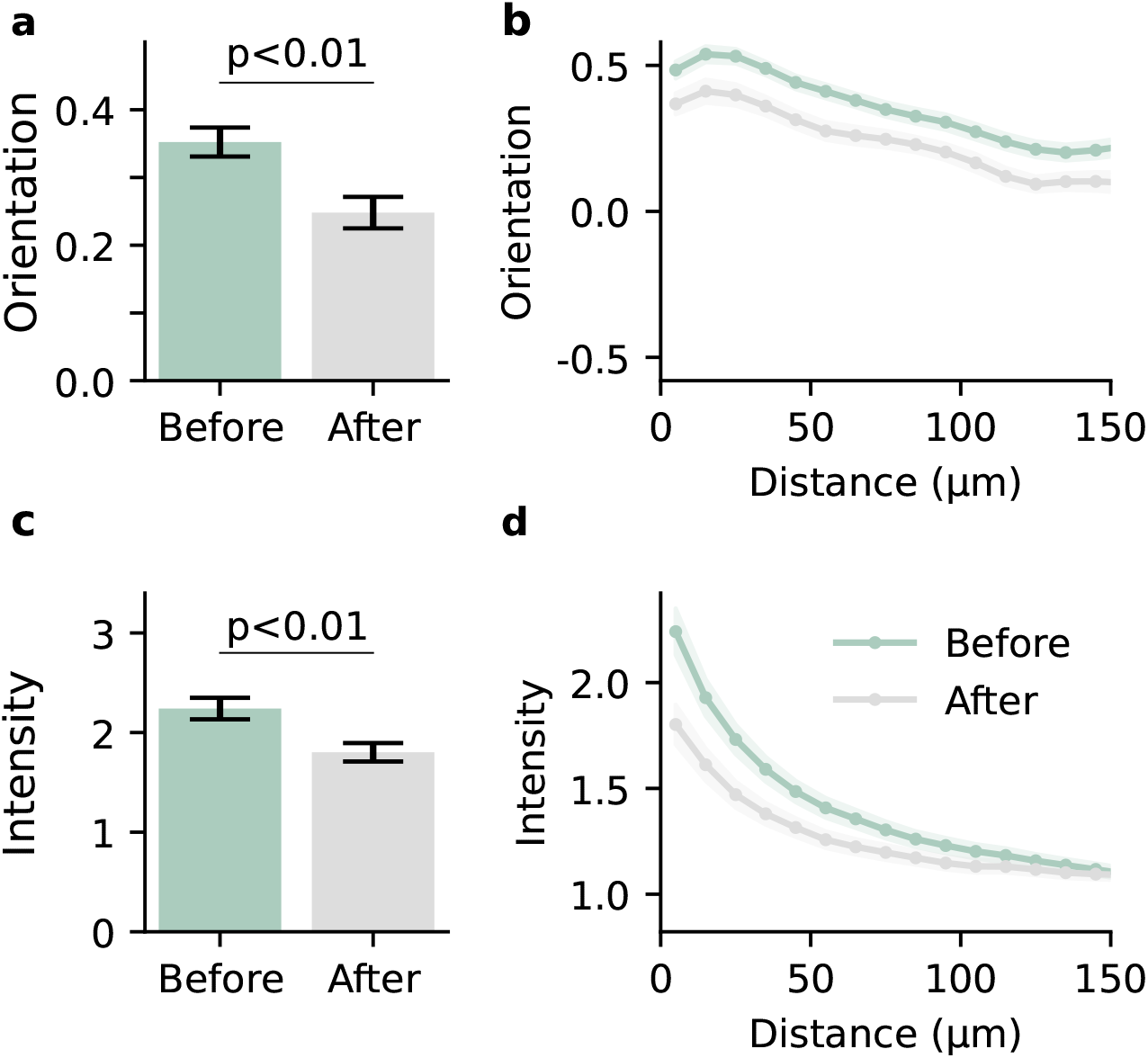
Non-elastic matrix remodelling around hepatic stellate cells. Averaged collagen orientation (**a**,) and collagen intensity c around hepatic stellate cells cultured for two days in 1.2 mg/ml collagen gels (Matrix Bioscience, Mörlenbach) before and after treatment with 10 μM cytochalasin D. Bars indicate mean ± se of 46 individual cells. Statistical significance is calculated using the independent sample t-test. Orientation propagation (**b**) and intensity propagation (**d**) averaged within 10 μm distance shells around the same cells. Shaded areas indicate mean ± se. The quantification of the residual collagen fiber orientation and intensity after drug-induced force relaxation allows for the quantification of non-elastic matrix remodelling.

**Supplementary Information 8:**
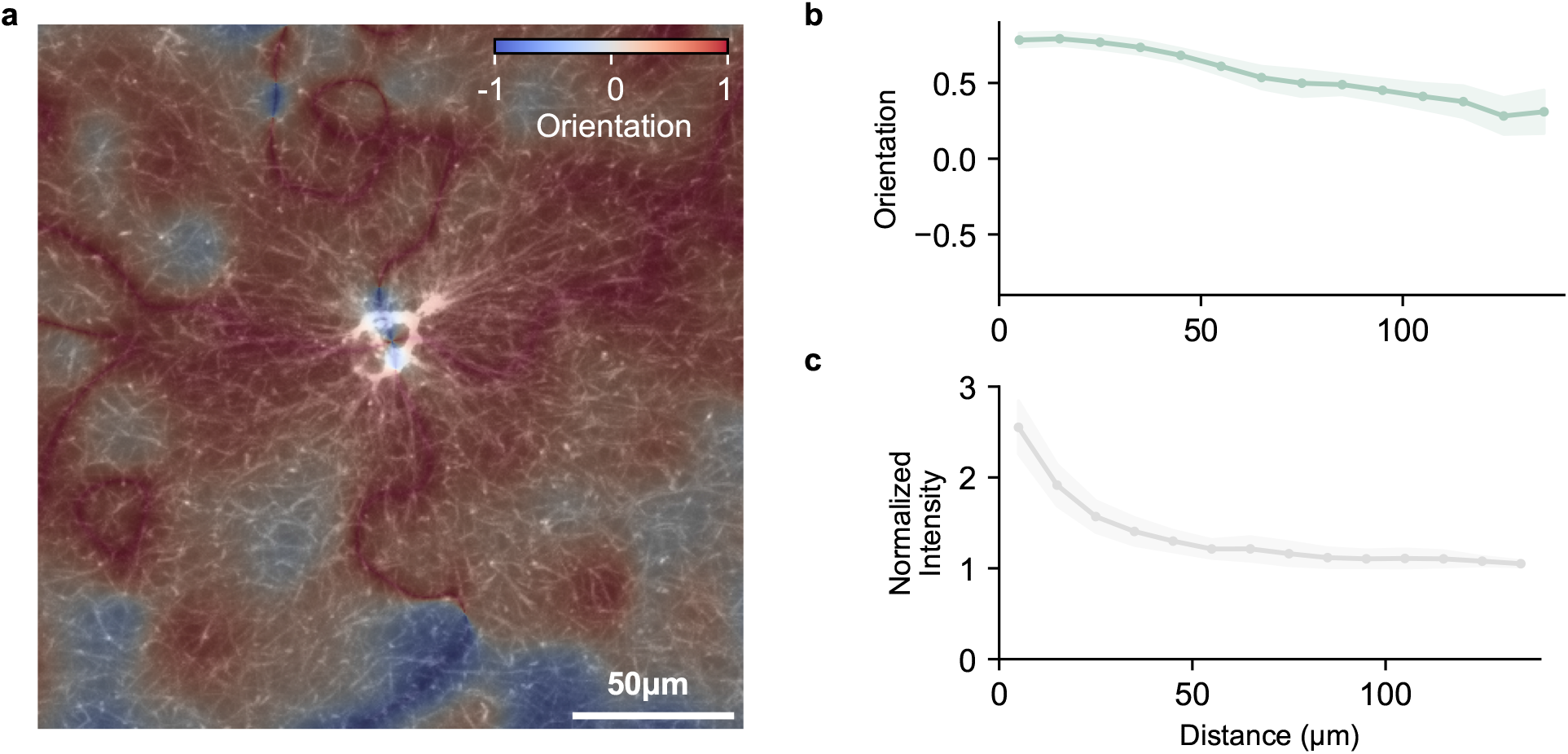
Fiber orientation and intensity in fibrin gels. **a**, Human lung fibroblasts are cultured for 4 hours in 3 mg/ml fibrin gels. Images show an overlay of the maximum intensity projected images (20 μm z-height around the cell) of the fluorescently labeled cell (Cell-TrackerGreen CMFDA, Thermo Fisher Scientific, Waltham) and the fluorescently labeled fibrin network (Alexa Fluor 647 NHS Succinimidyl Ester, Thermo Fisher Scientific, Waltham). Colormap indicates the fiber orientation calculated using a window-size of 10 μm. The orientation propagation (**b**) and the normalized fiber intensity (**c**) are calculated within 10 μm distance shells around the cells. Lines indicate the mean, and shaded areas indicate ± se for 6 individual cells.

**Supplementary Information 9:**
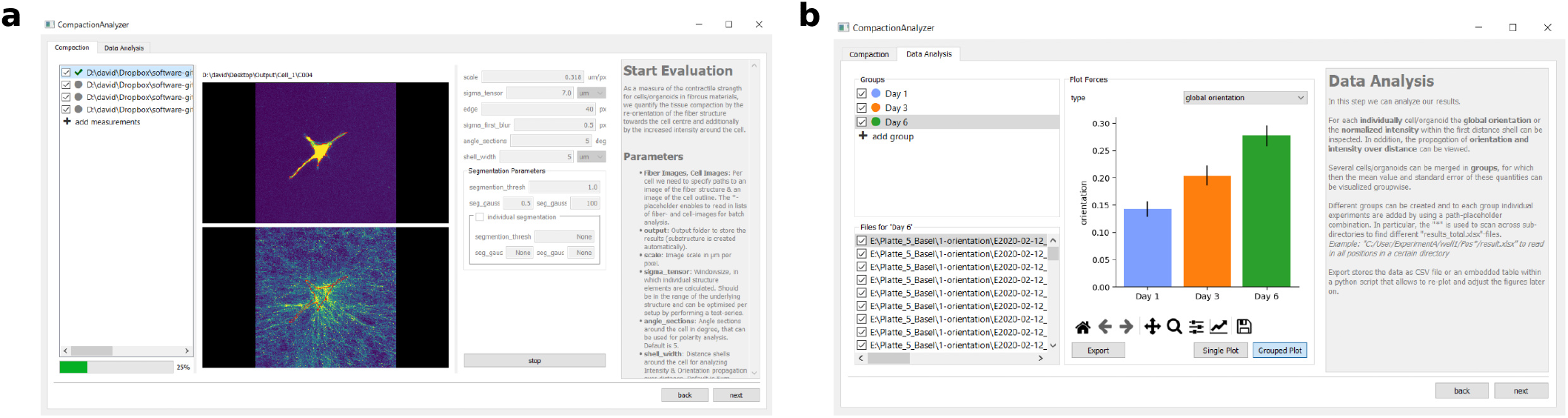
Graphical user interface (GUI) for the analysis of fiber orientation and intensity. Graphical user interface to measure fiber orientation and intensity around contractile cells embedded in fiber matrices. Pairs of fiber and cell images can be processed individually or batch-wise. **a**, Evaluation parameters for fiber detection and cell segmentation can be adjusted individually for each cell. **b**, Fiber orientation and intensity values can be evaluated individually or averaged over user-defined groups or treatment conditions. The software is available as an open-source Python package (29).

**Supplementary Information 10:**
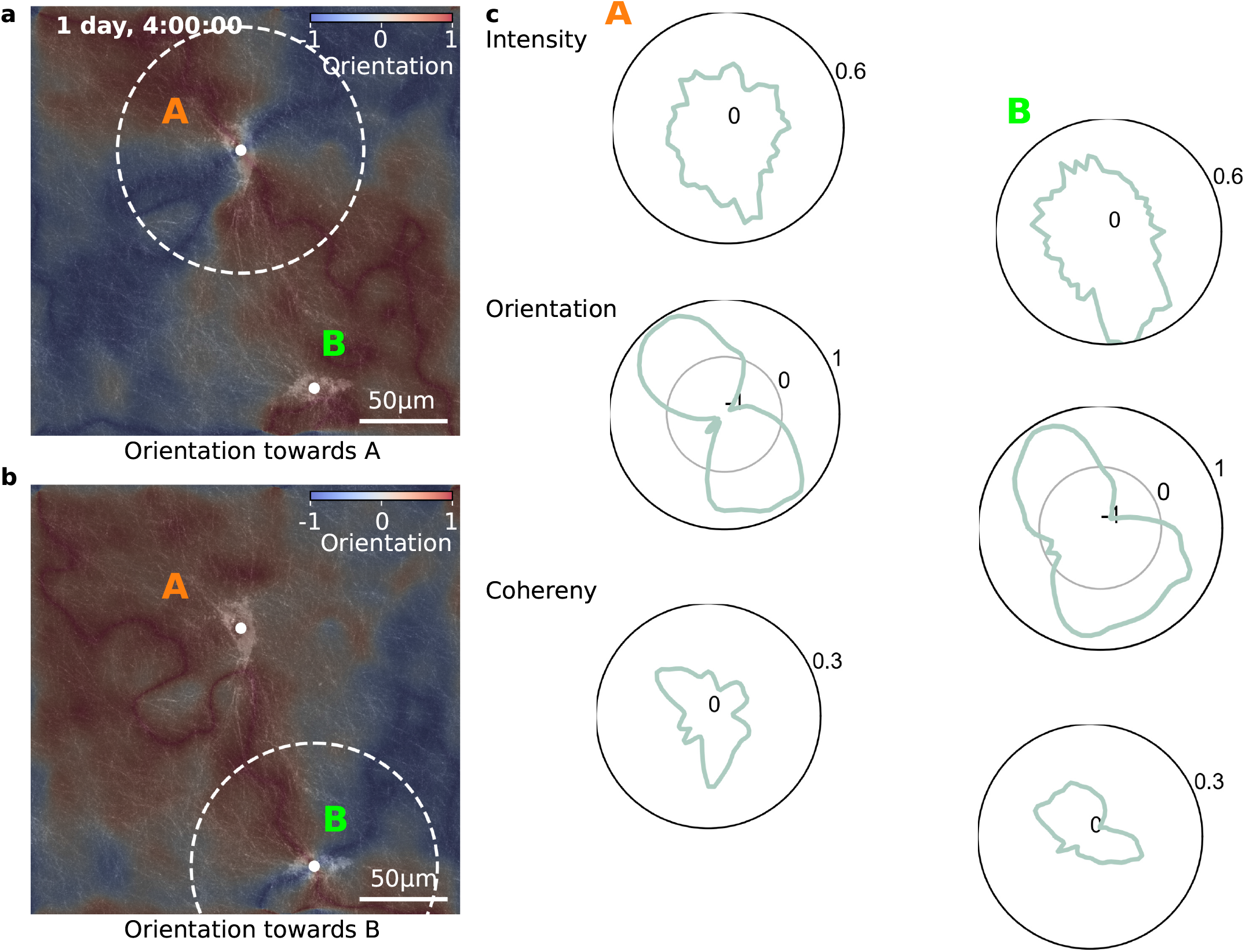
Mechanical cell-cell-interactions. Mechanical cell-cell-interactions between two hepatic stellate cells cultured in collagen gels for 28h with pronounced matrix fibers alignment between the cells. **a**,**b** Maximum intensity projected confocal reflection image (xyz=255×255×50 μm) of the collagen fibers around the cells overlaid with their orientation (color-coded in blue-red) towards cell A (**a**) and cell B (**b**). **c**, Polar plot of fiber intensity, fiber orientation, and fiber coherency around cell A (left) and cell B (right), averaged within a radius of 70 μm around the respective cells as indicated by the white dotted circles in **a**,**b**.

**Supplementary Information 11:**
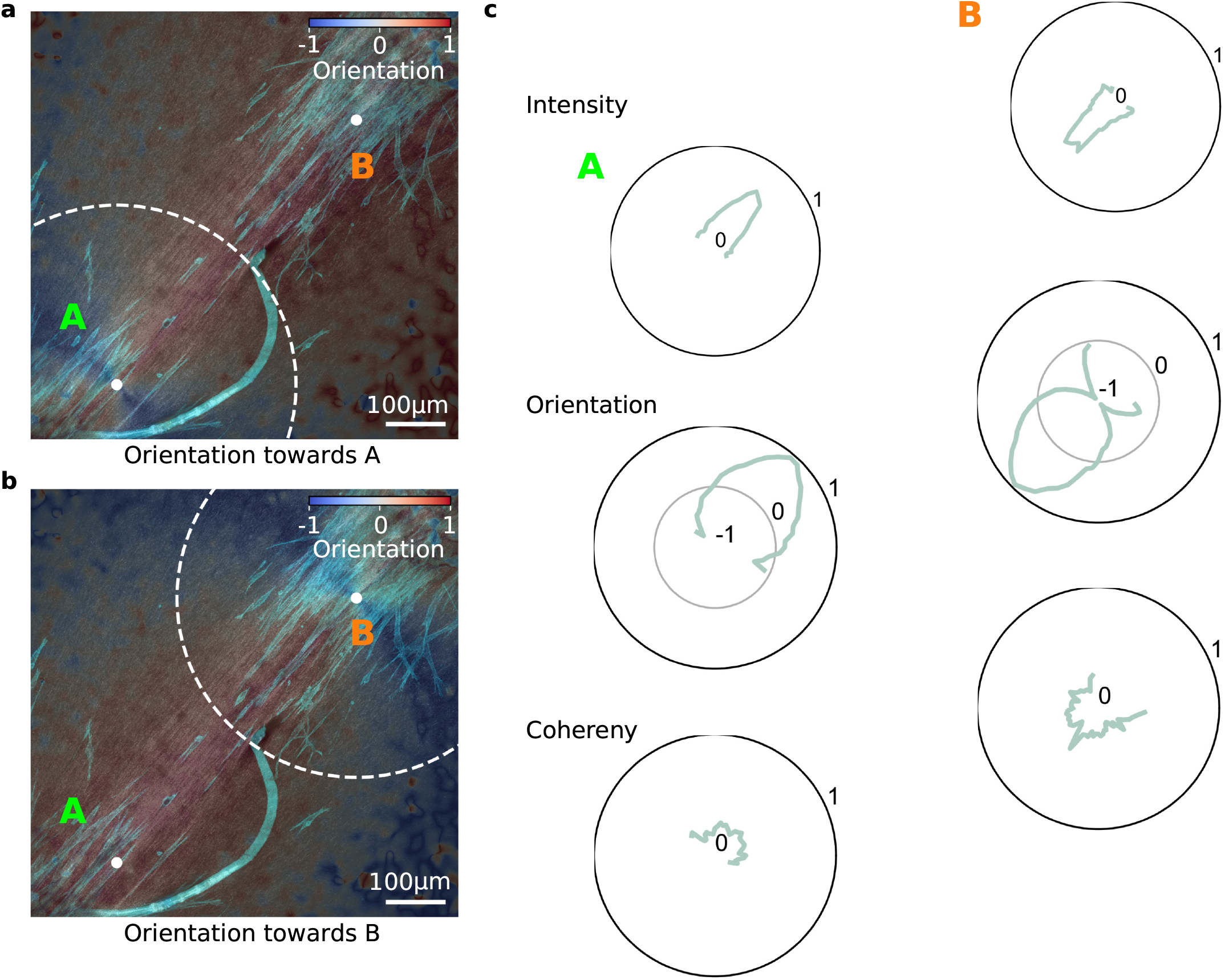
Collective cell-cell interactions. Collective cell-cell-interactions between two spheroids consisting of 4000 primary human cancer-associated fibroblasts embedded in 1.2 mg/ml collagen gels for 24h with pronounced matrix fibers alignment between the spheroids. **a**,**b** Maximum intensity projected confocal reflection image (738×738×8 μm) of the collagen fibers around the equatorial plane of the spheroids overlaid with the maximum intensity projected confocal fluorescence image (738×738×302 μm, green) of the actin stained cells (20 μg/ml Phalloidin-Tritc, Sigma-Aldrich, St. Louis). Orientation towards spheroid A (**a**) and spheroid B (**b**) are color-coded (blue-red). **c**, Polar plot of fiber intensity, fiber orientation, and fiber coherency around cell A (left) and cell B (right), averaged within a radius of 300 μm around the respective spheroid as indicated by the white dotted circles in **a**,**b**.

**Supplementary Information 12:**
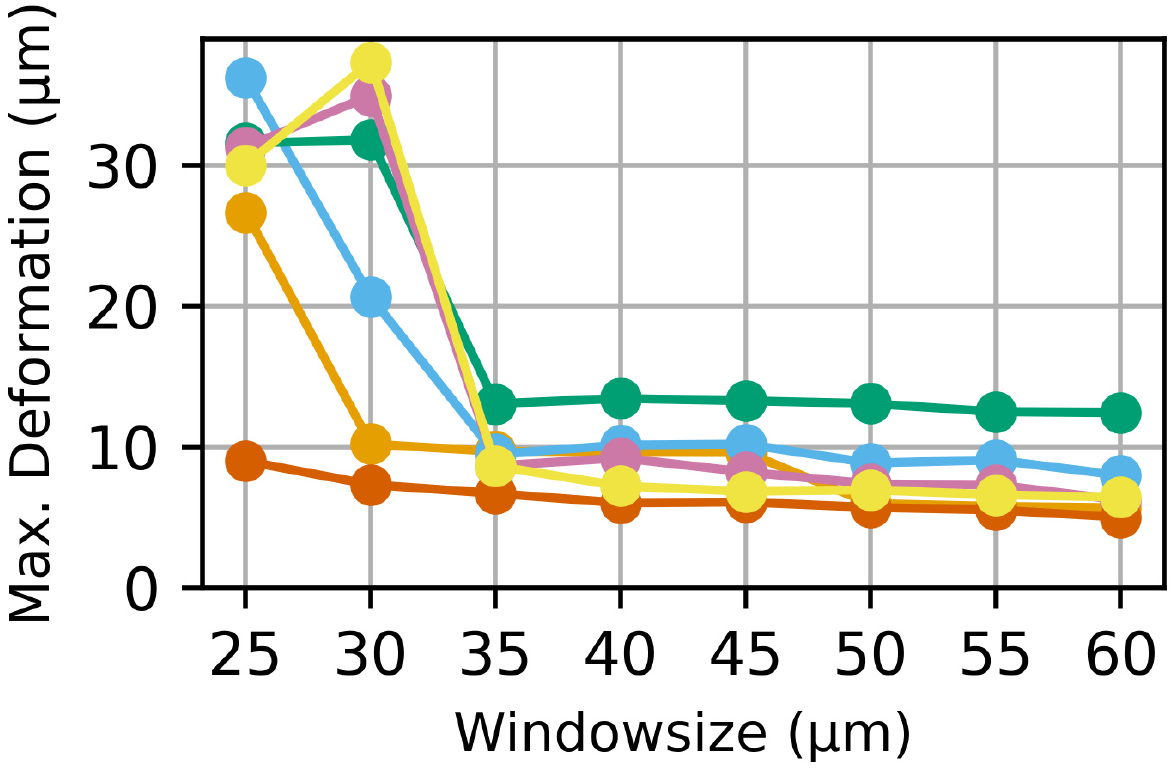
Window-size for 3D-PIV of hepatic stellate cells. Matrix deformations are determined around 6 different hepatic stellate cells (from Fig. 5f) with 3D particle image velocimetry (61) using different window-sizes (windows have an overlap of 60%, and the signal-to-noise filter is set to 1.3). The maximum measured deformations increase strongly for small window-sizes, but remain stable for larger window-sizes. The window-size of 35 μm was chosen as a compromise between overall resolution and noise.

**Supplementary Information 13: Video showing individual layers of an image stack of a hepatic stellate cell embedded in collagen**

Image stack (xyz=255×255×50 μm with a voxel-size of 0.32×0.32×2.5 μm) of a hepatic stellate cell embedded in a 1.0 mg/ml collagen gel acquired using second harmonic generation (collagen) and confocal fluorescence imaging (calcein stainded cell in orange).

**Supplementary Information 14: Video showing individual layers of an image stack of a lung fibroblast embedded in fibrin**

Image stack (xyz=256×256×58 μm with a voxel-size of 0.5×0.5×0.5 μm) of a lung fibroblast embedded in a 3.0 mg/ml fibrin gel acquired using fluorescence confocal microscopy.

**Supplementary Information 15: Video showing relaxation of cell forces using cytochalasin D**

Hepatic stellate cells are embedded in 1.2 mg/ml collagen gels (Matrix Bioscience, Mörlenbach) and cultured without (top) and with the addition of 10 μM Rock-inhibitor Y-27632 (bottom) for 2 days. The force-free reference state of the collagen is imaged 30 minutes after treatment with 10 μM cytochalasin D (Sigma-Aldrich, St. Louis). Images show maximum intensity projected confocal reflection image stacks (10 μm z-height) before and after cytochalasin D treatment. Scalebar indicates 50 μ m. Cells treated with Rock-inhibitor show reduced collagen contractions.

**Supplementary Information 16: Timelapse video showing 3D rendered hepatic stellate cell in collagen**

3D rendered representation of the hepatic stellate cell (green) shown in SI Video. 13 embedded in 1.0 mg/ml collagen (brown). Image stacks are acquired over time (dt≈30 min) starting after 7 hours culture in collagen. Cell is segmented using Yen thresholding (67). Collagen fiber images are Sato-filtered for ridge detection, highlighting the fiber structure (68). The transparency of the cell- and collagen image-stacks are determined by the intensity values according to a sigmoidal transfer function.

**Supplementary Information 17: Timelapse video showing 3D rendered hepatic stellate cells in collagen**

Same as in SI Fig. 16, but for two neighbouring hepatic stellate cells.

**Supplementary Information 18: Timelapse video showing maximum intensity projected images of hepatic stellate cells in collagen**

Maximum intensity projected images of the hepatic stellate cells shown in SI Video 16. Hepatic stellate cells are shown in blue and collagen is shown in brown.

**Supplementary Information 19: Timelapse video showing maximum intensity projected images of hepatic stellate cells in collagen**

Maximum intensity projected images of the hepatic stellate cells shown in SI Video 17. Hepatic stellate cells are shown in blue and collagen is shown in brown.

