## Supplementary figures and images for "Fiber alignment in 3D collagen networks as a biophysical marker for cell contractility"

### SI_Video_14

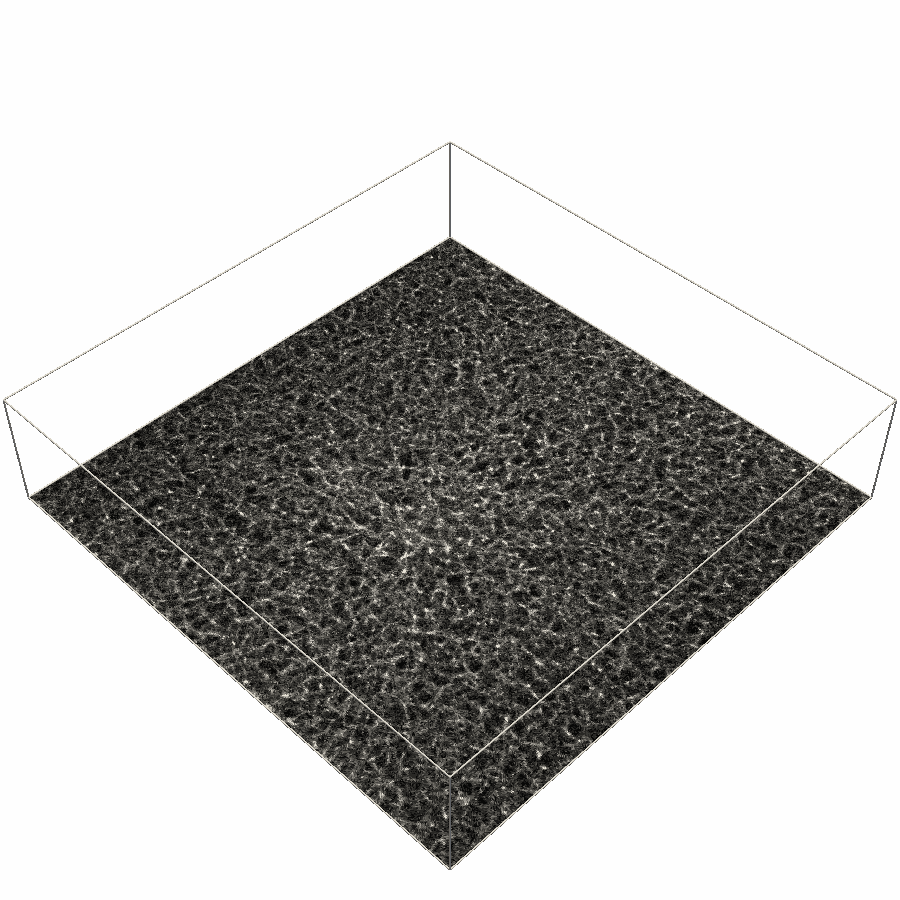

### SI_Video_15

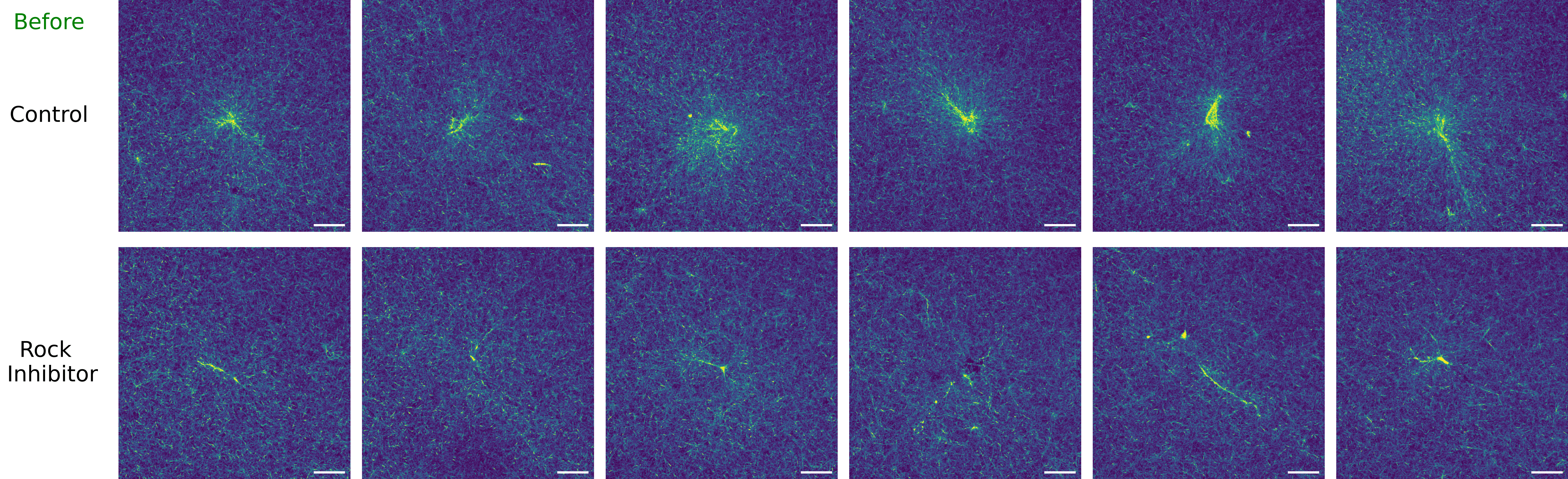
